# Novel hydroxamic acid derivative induces apoptosis and constrains autophagy in leukemic cells

**DOI:** 10.1101/2023.03.09.531973

**Authors:** Marten A. Fischer, Al-Hassan M. Mustafa, Kristin Hausmann, Ramy Ashry, Anita G. Kansy, Magdalena C. Liebl, Christina Brachetti, Andrea Piée-Staffa, Matthes Zessin, Hany S. Ibrahim, Thomas G. Hofmann, Mike Schutkowski, Wolfgang Sippl, Oliver H. Krämer

## Abstract

**Introduction:** Posttranslational modification of proteins by reversible acetylation regulates key biological processes. Histone deacetylases (HDACs) catalyze protein deacetylation and are frequently dysregulated in tumors. This has spurred the development of HDAC inhibitors (HDACi). Such epigenetic drugs modulate protein acetylation, eliminate tumor cells, and are approved for the treatment of blood cancers.

**Objectives:** We aimed to identify novel, nanomolar HDACi with increased potency over existing agents and selectivity for the cancer-relevant class I HDACs (HDAC1/-2/-3/-8). Moreover, we wanted to define how such drugs control the apoptosis-autophagy interplay. As test systems, we used human leukemic cells and embryonic kidney-derived cells.

**Methods:** We synthesized novel pyrimidine-hydroxamic acid HDACi (KH9/KH16/KH29) and performed in vitro activity assays and molecular modeling of their direct binding to HDACs. We analyzed how these HDACi affect leukemic cell fate, acetylation, and protein expression with flow cytometry and immunoblot. The publicly available DepMap database of CRISPR-Cas9 screenings was used to determine sensitivity factors across human leukemic cells.

**Results:** Novel HDACi show nanomolar activity against class I HDACs. These agents are superior to the clinically used hydroxamic acid HDACi vorinostat. Within the KH-series of compounds, KH16 (yanostat) is the most effective inhibitor of HDAC3 (IC_50_ = 6 nM) and the most potent inducer of apoptosis (IC_50_ = 110 nM; p<0.0001) in leukemic cells. KH16 though spares embryonic kidney-derived cells. Global data analyses of knockout screenings verify that HDAC3 is a dependency factor in human blood cancer cells of different lineages, independent of mutations in the tumor suppressor p53. KH16 alters pro- and anti-apoptotic protein expression, stalls cell cycle progression, and induces a caspase-dependent processing of the autophagy proteins ULK1 and p62.

**Conclusion:** These data reveal that HDACs are required to stabilize autophagy proteins through a suppression of apoptosis in leukemic cells. HDAC3 appears as a valid anti-cancer target for pharmacological intervention.

**Highlights:**

- Novel HDACi with nanomolar activity against leukemic cells were synthesized.
- HDACi of the KH-series are superior to a clinical grade HDACi.
- HDACi of the KH-series modulate acetylation and phosphorylation of proteins.
- The new HDACi KH16 regulates cell cycle arrest, apoptosis, and autophagy.
- Apoptosis acts upstream of autophagy in KH16-treated cells.

## Introduction

Histone deacetylases (HDACs) are Zn^2+^- or NAD^+^-dependent enzymes that control the acetylation status of cellular proteins in procaryotes and eucaryotes. 18 mammalian HDACs are classified into four classes, based on their homology to yeast HDACs. Class I HDACs comprise HDAC1,-2,-3,-8, class II is divided into IIA being HDAC4,-5,-7,-9 and class IIB being HDAC6,-10, and class IV contains only HDAC11, which has properties of classes I and II [1, 2]. The HDAC classes I, II, and IV use Zn^2+^ as a cofactor and the class III HDACs SIRT1-7 utilize NAD^+^ as a cofactor [2, 3].

Cancerous tissues frequently overexpress class I HDACs and have pathology-linked altered acetylation patterns compared to healthy tissues [1,4,5]. Such findings have stimulated the development of HDAC inhibitors (HDACi). HDACi against Zn^2+^-dependent HDACs have been approved by the U.S. food-and-drug administration (FDA) for the therapy of human blood tumors [1,4,5]. This was preceded by the use of weakly active, millimolar fatty acid HDACi in compassionate protocols in the last century [2]. The micromolar hydroxamic acid vorinostat (SAHA) was the first clinically approved HDACi. The hydroxamic acid belinostat, as well as the depsipeptide romidepsin are additional HDACi that are approved by the U.S. FDA. The China FDA approved the HDACi tucidinostat, also called chidamide, for clinical use against subtypes of leukemia [1,4,5]. Hydroxamic acids are pan-HDACi with a variable activity against all Zn^2+^-dependent HDACs, romidepsin specifically inhibits class I HDACs, and the benzamide tucidinostat blocks HDAC1-3 and HDAC10 [6, 7]. Additional HDACi, such as the pan-HDACi abexinostat and pracinostat have produced promising results in clinical trials [8, 9]. Therefore, synthesizing and characterizing HDACi that are based on the hydroxamic acid scaffold is a promising approach to identify more effective epigenetic drug candidates for cancer treatment.

A common feature of clinical grade HDACi is a nano- to micromolar inhibition of class I HDACs. This encourages the synthesis of HDACi with selectivity for these enzymes. Beyond high activity against tumor cells, selective HDACi are expected to have less side-effects than pan-HDACi [2]. HDACi with the benzamide group as a molecular warhead to complex Zn^2+^ can have remarkable class I HDAC selectivity but lack the low nanomolar activities of depsipeptides and hydroxamic acids [6,7,10,11]. There is an intense search for hydroxamic acids with a very potent inhibition of individual class I HDACs.

The best-characterized anti-proliferative effects of HDACi include the induction of cell cycle arrest and non-inflammatory cell death through apoptosis. The molecular mechanisms underlying the induction of cell cycle arrest by HDACi involve an increase in cyclin-dependent kinase inhibitors and a downregulation of cyclin-dependent kinases. HDACi-induced apoptosis relies on an upregulation of pro-apoptotic and a downregulation of anti-apoptotic proteins, including members of the BCL2 family and a disruption of kinase signaling cascades. These effects can occur through RNA- and protein-dependent mechanisms [1,4,5,12]. HDACi additionally halt tumor cell proliferation through the induction of DNA replication stress and DNA damage [13, 14]. Recent data show that HDACi also modulate a “self-eating” mechanism of cells termed autophagy. This pathway digests cellular proteins and organelles under stress to promote cell survival [15, 16]. Accordingly, the induction of autophagy frequently antagonizes pro-apoptotic properties of HDACi [5,17-19].

A nanomolar HDACi with class I HDAC selectivity is both a clinically interesting molecule and a pharmacological tool to assess the specific functions of HDACs. We developed and characterized novel pyrimidine-hydroxamic acids regarding their potency against leukemic cells. We compared these compounds with established HDACi and analyzed the impact of the most potent new HDACi on key proteins controlling apoptosis and autophagy. As testbeds, we used human leukemic cells and kidney-derived cells, and we analyzed global datasets. The most promising compound, KH16 (yanostat), preferentially inhibits HDAC3, efficiently induces apoptosis, and disables autophagy through caspase activation.

## Materials and methods

### HDACi

SAHA was purchased from Selleck Chemicals, Munich, Germany. KH9, KH16 (yanostat), and KH29 were synthesized as mentioned in the supplemental information file **SI** and Figs **S1**-**S12**. All drugs were kept as stock solutions in DMSO at -80°C and diluted in PBS immediately before treatment.

### In vitro HDAC enzymatic assay

Recombinant human HDAC1, HDAC2, HDAC3/NCOR1, and HDAC6 were purchased from ENZO Life Sciences AG (Lausen, CH). Recombinant human HDAC11 was produced by Barinka et al. [20]. Recombinant human HDAC8 was produced by Romier et al. [21]. The in vitro tests with recombinant HDAC1-3 were performed with a fluorogenic peptide derived from p53 (Ac-RHKK(Acetyl)-AMC). The measurements were performed in assay buffer (50 mM HEPES, 150 mM NaCl, 5 mM MgCl_2_, 1 mM TCEP and 0.2 mg/mL BSA, pH 7.4 adjusted with NaOH) at 37°C. Inhibitors at different concentrations were incubated with 10 nM HDAC1, 3 nM HDAC2, or 3 nM HDAC3 (final concentration) for at least 5 min. The reaction was started with the addition of the fluorogenic substrate (20 µM final concentration) and incubated for 30 min for HDAC2 and HDAC3 and 90 min for HDAC1. The reaction was stopped with a solution of 1 mg/mL trypsin and 20 µM SAHA in 1 mM HCl and incubated for 1 h at 37°C. The fluorescence intensity was recorded with an Envision 2104 Multilabel Plate Reader (PerkinElmer, Waltham, MA), with an excitation wavelength of 380 ± 8 nm and an emission wavelength of 430 ± 8 nm. The received fluorescence intensities were normalized with uninhibited reaction as 100% and the reaction without enzyme as 0%. A nonlinear regression analysis was done to determine IC_50_ values. Dose-response-curves for HDAC6 were determined as described [20], with the substrate (Abz-SRGGK(thio-TFA)FFRR-NH2). The substrate concentration was 50 µM and the enzyme concentration was 10 nM for HDAC6. The HDAC11 inhibition assay was performed as described before [22]. The fluorescence intensity was recorded with the above-mentioned reader, with an excitation wavelength of 330 ± 75 nm and an emission wavelength of 430 ± 8 nm. The enzyme inhibition of HDAC8 was determined by using a homogenous fluorescence assay [21]. The enzyme was incubated for 90 min at 37°C, with the fluorogenic substrate ZMAL (Z(Ac)Lys-AMC) in a concentration of 10.5 μM and increasing concentrations of inhibitors. Fluorescence intensity was measured at an excitation wavelength of 390 nm and an emission wavelength of 460 nm in a microtiter plate reader (BMG Polarstar).

### Docking study

Protein structures were retrieved from the Protein Data bank (PDB) (HDAC1 PDB ID: 4BKX, HDAC2 PDB ID: 5IWG. HDAC3 PDB ID: 4A69) [23–25]. Protein structures were prepared using the Protein Preparation Wizard module in Schrödinger Suite [26]. Hydrogen atoms and missing side chains were added. Water molecules involved in protein-ligand interaction in HDAC1, HDAC2, and HDAC3, were kept for the docking study. The protonation states and tautomeric forms of the amino acids were optimized using PROPKA tool at pH 7.0. The potential energy of the three optimized structures was minimized using OPLS3e force-field [27]. Ligands were prepared using the LigPrep module in Schrödinger Suite using OPLS3e force-field. KH compounds were docked in the deprotonated hydroxamate and protonated piperazine forms. Conformations of prepared ligands were generated using the Confgen tool in Schrödinger Suite by applying 64 conformers per each ligand and minimizing the conformers. Molecular docking studies were conducted by applying the Glide program in Schrödinger Suite. The grid box was generated with 10*10*10 Å size using the Receptor Grid Generation module in Schödinger19. Standard Precision (SP) mode with flexible ligand sampling was utilized for docking. To validate the docking protocol, re-docking studies were done using the co-crystallized ligands. Root mean square deviation (RMSD) values for the docked inhibitors were in all cases below 0.5 Å as already reported [10]. Docking poses were visualized in the MOE2018.01 program [28].

### Cell culture

Experiments were carried out with HEL cells (erythroblast cell line from the bone marrow of a 30-year-old, White, male patient with erythroleukemia), Ramos cells (B-lymphocyte cell line from a 3-year-old, White, male Burkitt’s Lymphoma patient), RS4-11 cells (lymphoblast cells from the marrow of a 32-year-old, White, female patient with acute lymphoblastic leukemia), and human embryonic kidney cells (HEK293T cells, originally from a female fetus). The cells were kind gifts from Dr. Grez (Frankfurt/Main, Germany), Dr. Kosan (Jena, Germany), Prof. Böhmer (Jena, Germany), Prof. Heinzel (Jena, Germany), and were originally from the DSMZ (Braunschweig, Germany). Cells were cultured under sterile conditions at 37°C and 5% CO_2_ in Roswell Park Memorial Institute-1640 (RPMI1640) (Sigma-Aldrich) for leukemic cells or Dulbecco’s Modified Eagle Medium (DMEM) (Sigma-Aldrich) for HEK293T cells. Media were supplemented with 5% fetal calf serum (FCS) (Sigma-Aldrich) and 1% penicillin/streptomycin (Gibco). As previously reported, using 5% instead of 10% FCS reduces cell stress during proliferation [29]. Cell density was kept below 400,000 cells per mL. Cell suspensions were centrifuged (300 x g, 5 min), medium was discarded, and replaced by fresh medium. 200,000 cells per mL were seeded per well and placed back in the incubator for at least two hours prior to any treatment.

### Immunoblot and antibodies

Immunoblot was done as recently described by us [30–32].

Primary antibodies used in this work:

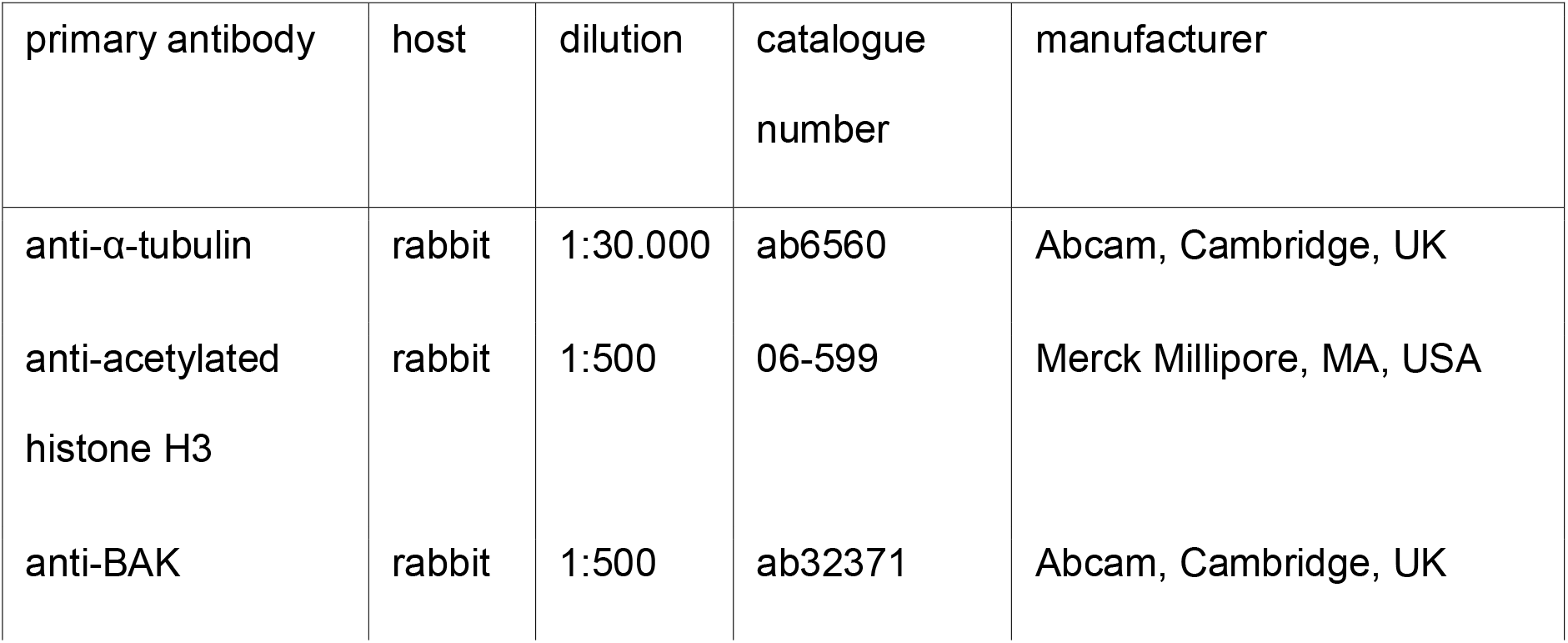
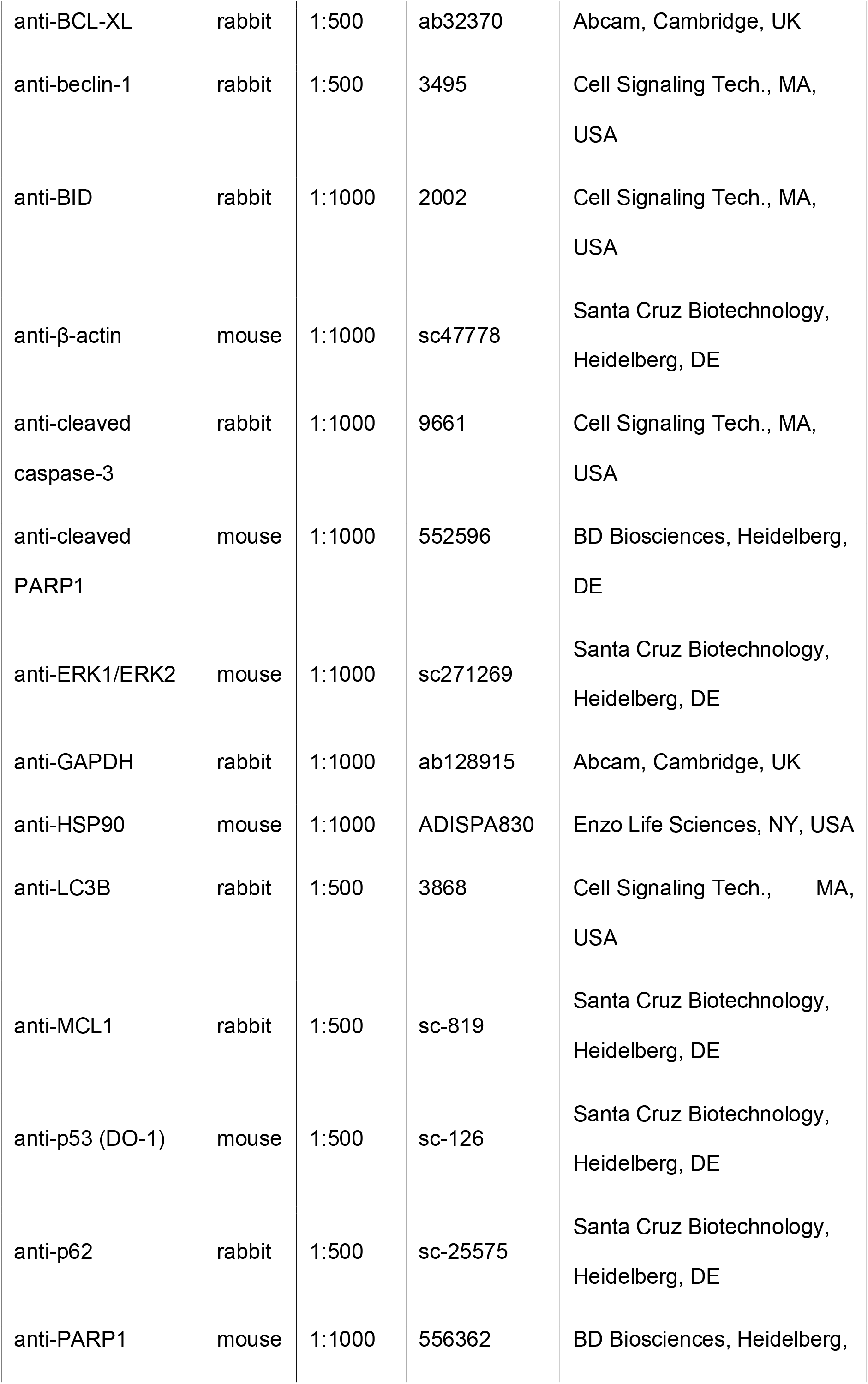
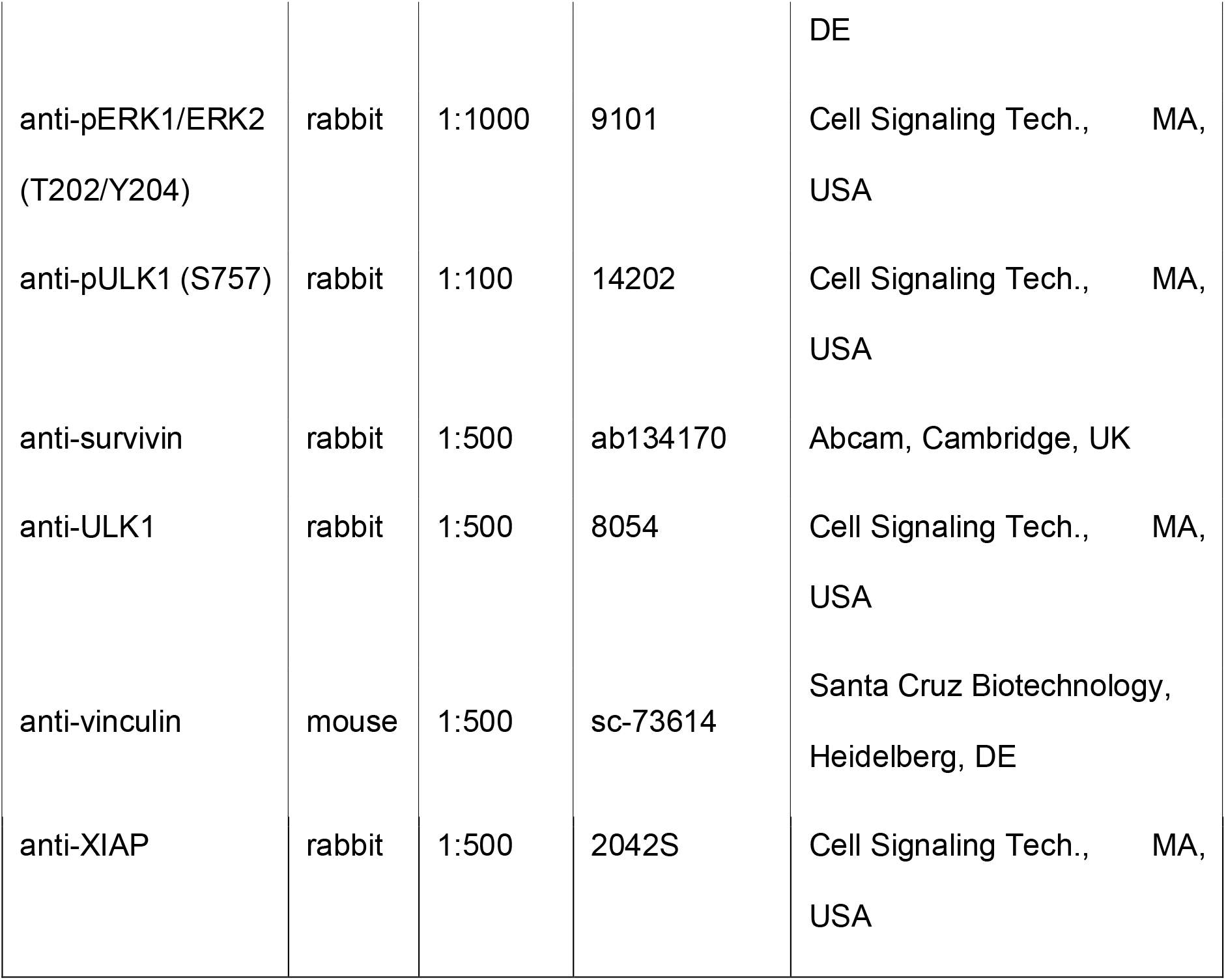

Secondary antibodies used in this work:

| secondary antibody | host | catalogue<br>number | manufacturer |
| --- | --- | --- | --- |
| IRDye®680RD anti-mouse IgG | goat | 925-68070 | LI-COR, NE, USA |
| IRDye®680RD anti-rabbit IgG | goat | 925-68071 | LI-COR, NE, USA |
| IRDye®800CW anti-mouse IgG | goat | 925-32210 | LI-COR, NE, USA |
| IRDye®800CW<br>anti-rabbit IgG | goat | 925-32211 | LI-COR, NE, USA |
| anti-mouse IgG, HRP-linked<br>antibody | horse | 7076 | Cell Signaling Tech.,<br>MA, USA |
| anti-rabbit IgG, HRP-linked<br>antibody | horse | 7074 | Cell Signaling Tech.,<br>MA, USA |

### Annexin-V/propidium iodide (PI) staining

This assay was previously described by us [30–32]. Annexin-V binds to phosphatidylserine, which becomes exposed on the outer membrane of early and late apoptotic cells. PI stains DNA and is utilized as a marker of late-stage apoptosis and necrosis, with non-intact cell membranes. In brief, cells were seeded at 200.000 cells per mL in 24-well plates and harvested into flow cytometry tubes. Note that HEL cell cultures have some cells attached to the culture dishes. Such cells should be collected by trypsinization. Cells were centrifuged (300 x g, 5 min), washed twice in PBS, resuspended in 2.5 µL annexin-V (BD Bioscience) diluted in 50 µL annexin-V binding buffer (1 mM HEPES, 0.14 M NaCl, 0.25 mM CaCl_2_, adjusted to pH 7.4 with KOH), and kept in the dark. After 15 min, 10 µL of PI (Sigma-Aldrich) solution (50 µg/mL, diluted in 430 µL annexin-V binding buffer) were added. Cells were analyzed after 10 min with a FACS Canto II flow cytometer. Data were examined with the FACS Diva software. To calculate IC_50_ values, we chose logarithmic scale concentrations from 2.5 nM to 5 µM in Graphpad Prims 6.

### Cell cycle analysis

We recorded cell cycle profiles by flow cytometry [30–32]. PI is a marker for DNA contents in different phases of the cell cycle. From G1 phase to G2/M phase, the cellular DNA content increases from 2N to 4N. Apoptotic cells have DNA contents below 2N (termed subG1 fractions). In brief, cells were seeded at 200.000 cells per mL in 24-well plates. Both floating and adherent cells were harvested and washed twice with ice-cold PBS. After discarding the PBS, 1 mL ice-cold 80% EtOH was added dropwise while vortexing to fix the cells. After storage at 20°C for at least 2 h, samples were centrifuged, and ethanol was discarded. To prevent false measurement of RNA instead of DNA, fixed cells were treated with 333 µL PBS supplemented with 1 µL RNAse (10 mg/mL) (Roche) at 37°C for 30 min. Cells were treated with 164 µL PI-staining solution (50 μg/mL) and measured after 10 min using a FACS Canto II. Further data analysis was carried out with FACS Diva and Prism 6.

### Cancer dependency map (DepMap) data base analysis

CRISPR dependency data (CHRONOS scores) and *TP53* (gene encoding the transcription factor p53) mutation data for cell lines of myeloid and lymphoid lineages were both downloaded from the publicly available 2022 Q4 DepMap release using the Broad Institute’s DepMap portal (https://depmap.org/portal/). The Wilcoxon rank-sum test, which does not require that the data show a normal distribution test, was applied to compare dependency scores between *TP53* wild-type and *TP53* mutant cell lines.

Software used in this work

| Software | Developers |
| --- | --- |
| BD FACSDiva™ CS 6 | BD Biosciences, Heidelberg, DE |
| Cell Imaging Software | Olympus Soft Imaging Solutions, Münster, DE |
| GraphPad Prism Version 6 | GraphPad Software, CA, USA |
| Image Studio Lite 5.2 | LI-COR, NE, USA |
| MS Office 365 | Microsoft, Unterschleißheim, DE |
| Odyssey Version 3.0 | LI-COR, NE, USA |

### Statistical analysis

Statistical analyses were done using two-tailed Student’s t test with Microsoft Excel (\**p*<0.05, \*\**p*<0.01, \*\*\**p*<0.001). All values are displayed as mean ± SEM.

Statistical analysis of Flow cytometry Data was done using two-way ANOVA tests with Bonferroni correction using GraphPad Prism 6 (*p<0.5, **p<0.01, ***p<0.001, ****p<0.0001)

## Results

### In vitro activity of new HDACi of the KH-series: KH9, KH16, and KH29

We synthesized a battery of pyrimidine-based hydroxamic acids as potential HDACi and tested them with an established enzymatic assay [10] in vitro (KH-series, **Table 1**). Synthesis details and analytical characterization of these compounds are described in the supplementary information file **SI** and Figs. S1**-S12**. KH9, KH16, and KH29 have low nanomolar in vitro activity against class I HDACs (**Table 1**). The three compounds were similarly active against HDAC2 (IC_50_ = 22-34 nM). KH9 and KH29 were most active against HDAC1 (IC_50_ = 9 nM) and KH16 was most active against HDAC3 (IC_50_ = 6 nM). Except for KH29, the KH-compounds were less active against the class I HDAC HDAC8 (IC_50_ = 49-530 nM) and the class IIB HDAC HDAC6 (IC_50_ = 25-460 nM). All agents showed far weaker activity against the class IV HDAC HDAC11 (IC_50_ = 870-1500 nM) (**Table 1**).

KH9, KH16, and KH29 contain the hydroxamic acid moiety which is present in the clinically approved HDACi SAHA. SAHA is an at least eightfold less potent inhibitor of HDAC1, HDAC2, and HDAC3 than KH9, KH16, and KH29. SAHA and KH29 are similarly effective against HDAC6, and both are at least 10-fold more potent inhibitor of HDAC6 than KH9 and KH16 are (**Table 1**).

These data show that the KH-compounds are low nanomolar inhibitors of the cancer-associated enzymes HDAC1, HDAC2, and HDAC3 in vitro.

### Assessment of apoptosis induction by KH compounds

The low nanomolar inhibitory activity of these agents encouraged us to test them in leukemic cells. Since HDACi are considered for the treatment of myeloproliferative neoplasms (MPNs) [2, 33], we chose HEL erythroleukemia cells as testbed to analyze cellular effects of our new HDACi. We treated these cells with KH9, KH16, and KH29 and assessed early apoptosis (annexin-V positivity) and late apoptosis (annexin-V/PI positivity) by flow cytometry. We calculated the IC_50_ values for total apoptosis induction by the KH compounds and used the clinically validated HDACi SAHA for comparison. KH9 induced apoptosis starting at approximately 50 nM and with a corresponding IC_50_ value of 164.8 ± 2.13 nM. KH29 was similarly effective, with apoptosis induction starting at 55 nM and an IC_50_ value of 162.0 ± 2.12 nM. KH16 was the most effective, starting to trigger apoptosis at 30 nM and with an IC_50_ value of 110.0 ± 2.10 nM. SAHA evoked apoptosis at a concentration starting from approximately 400 nM and with an IC_50_ value of 1214.4 ± 1.92 nM (**Fig. 1A-D**). KH16 had the lowest IC_50_ value of the KH-compounds and an 11-fold lower IC_50_ than that of SAHA. Comparison of IC_50_ values demonstrated that KH9, KH16, and KH29 were significantly more effective than SAHA (p<0,0001) (**Fig. 1E**).

**Figure 1:**
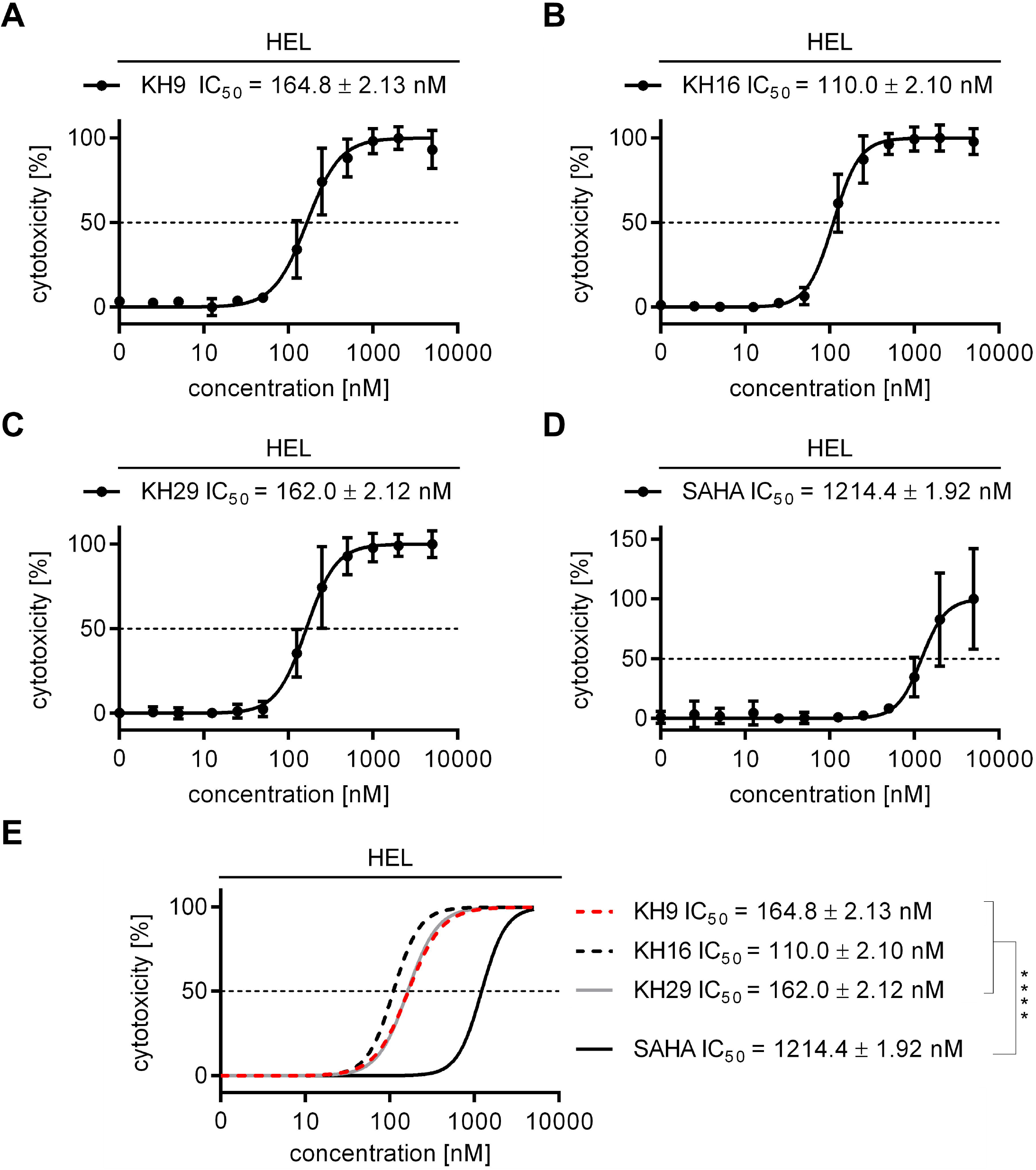
HDACi of the KH-series induce apoptosis of leukemic cells. (A-D) HEL cells were treated with increasing concentrations ranging from 25 nM to 5 µM of either KH9, KH16, KH29, or SAHA for 24 h. Their cytotoxicity was determined by annexin-V/PI staining and flow cytometry. (E) Overlay of data collected in (A-D). IC_50_ values were determined from independent experiments (KH-series n=4, SAHA n=3; p<0,001 for SAHA versus the KH-compounds).

These results show that the three KH compounds are up to 11-fold more potent than their parent compound, the FDA-approved HDACi SAHA, in leukemic cells. These data correspond to the notion that the KH-compounds have a higher in vitro activity against HDAC1, HDAC2, and HDAC3 than SAHA. KH16 exerts the strongest impact on HDAC3 in vitro and this translates into the strongest impact on the leukemic cell survival.

### KH16 bind class I HDACs through structurally defined regions

To rationalize the high inhibitory potency of the structurally highly similar KH16 and KH29 for HDAC1, HDAC2, and HDAC3 (**Table 1**), we docked them into available crystal structures. Modelling the interaction of KH-compounds with HDAC1, HDAC2, and HDAC3 unraveled that in all three HDACs, the hydroxamate coordinates the Zn^2+^ at the bottom of the catalytic clefts of the HDACs. The pyrimidine ring is sandwiched between two conserved Phe residues in the acetyllysine tunnel. The protonated piperazine forms a salt-bridge with the Asp99 residue in HDAC1/HDAC2 (corresponding to Asp92 in HDAC3). For HDAC1 and HDAC2, the capping group (methoxyphenyl of KH9, indole of KH16, and methylindole of KH29) makes van-der-Waals interaction with the Phe200 moiety in HDAC1/HDAC2 and the corresponding Phe200 residue in HDAC3. The overall binding mode of the inhibitors to HDAC1 and HDAC2 is highly similar. In case of HDAC3, KH16 has a different orientation of the capping group that is stabilized by a hydrogen bond between its indole NH and the Asp92 moiety. This interaction cannot be observed for the methylated derivatives KH29 and KH9, which do not possess a hydrogen bond donor in the capping group. This interaction is also not observed for KH16 docked to HDAC1 and HDAC2 due to a different conformation of this part of the binding pocket. Thus, this unique interaction profile can in part explain the high potency of KH16 for HDAC3 (**Fig 2A-C**).

**Figure 2:**
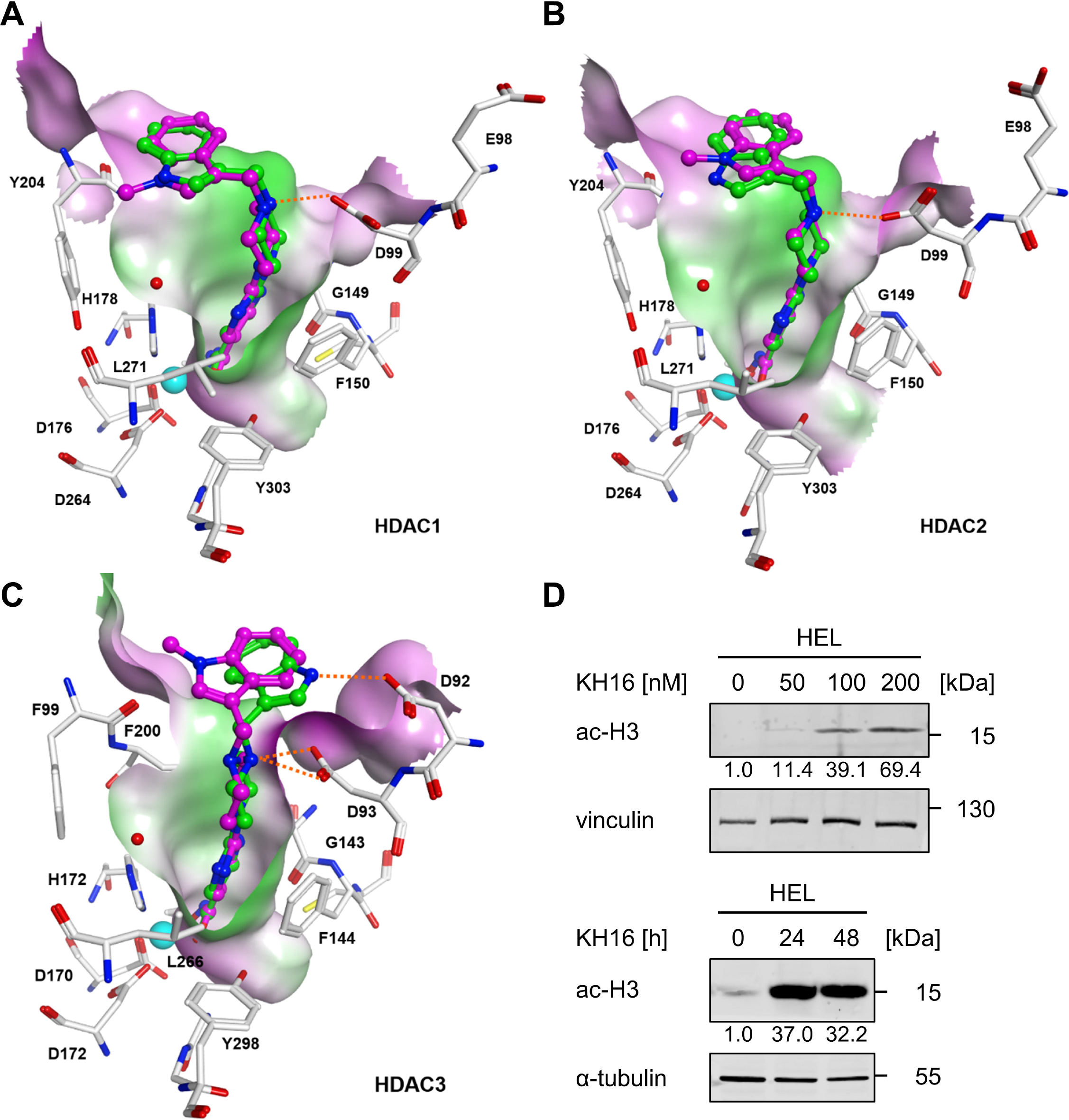
Binding mode of KH compounds to class I HDACs and accumulation of hyperacetylated histone H3. (A-C) Binding mode of KH16 (colored green) and KH29 (colored magenta) at HDAC1, HDAC2, and HDAC3. Hydrogen bonds and salt bridges between inhibitors and proteins are shown as orange-colored dashed lines. The molecular surfaces of the binding pockets are displayed and colored according to their hydrophobicities (green color indicates hydrophobic regions, magenta color indicates polar regions). (D) HEL cells were treated with 50, 100, or 200 nM KH16 for 24 h or 100 nM KH16 for 24 h to 48 h. Immunoblots were performed with antibodies against acetylated histone H3 and vinculin or α-tubulin as independent loading controls; ac-H3, acetylated histone H3. Data represent two or more independent repetitions. Numbers indicate densitometry values for the proteins, divided by those for the loading controls (Odyssey quantitative system). Untreated samples are set as 1.

Of the three KH compounds, KH16 is the most potent in vitro and in cells. To investigate if KH16 has on-target activity for the cancer-relevant class I HDACs, we analyzed the acetylation of their prototypical target histone H3 in cells [34]. We applied 50-200 nM KH16 for 24 h or 100 nM for 24 h to 48 h. KH16 increased acetylated histone H3 levels dose-dependently up to nearly 70-fold (**Fig. 2D**). 100 nM KH16 induced an over 30-fold increase in acetylated histone H3 at 24 h and this effect persisted over at least 48 h (**Fig. 2D**). These data illustrate the interaction of KH16 and KH29 with class I HDACs and verify that KH16 inhibits class I HDACs in leukemic cells.

### KH16 causes cell cycle arrest and DNA fragmentation

Of the KH-compounds KH16 has the lowest IC_50_ value (**Fig. 1E**). Therefore, we used KH16 for all further experiments with tumor cells. We used flow cytometry to test if the induction of protein hyperacetylation and apoptosis by KH16 are linked to cell cycle arrest of HEL cells. As expected, untreated HEL cells showed a typical cell cycle distribution in the G1, S, or G2/M phases, and only 7.6 ± 2.2% of cells were detectable as subG1 fractions with fragmented DNA (**Fig. 3A**). Treatment with 100 nM KH16 for 24 h decreased the numbers of cells in the S phase and G2/M phases. After 48 h, the G2/M phase population was decreased significantly by KH16. The numbers of cells in the G1 phase were not significantly changed by KH16 after 24 h and 48 h. The subG1 fractions significantly increased to 40 ± 11.2% and 55 ± 3.5% in a time-dependent manner (**Fig. 3A**).

**Figure 3:**
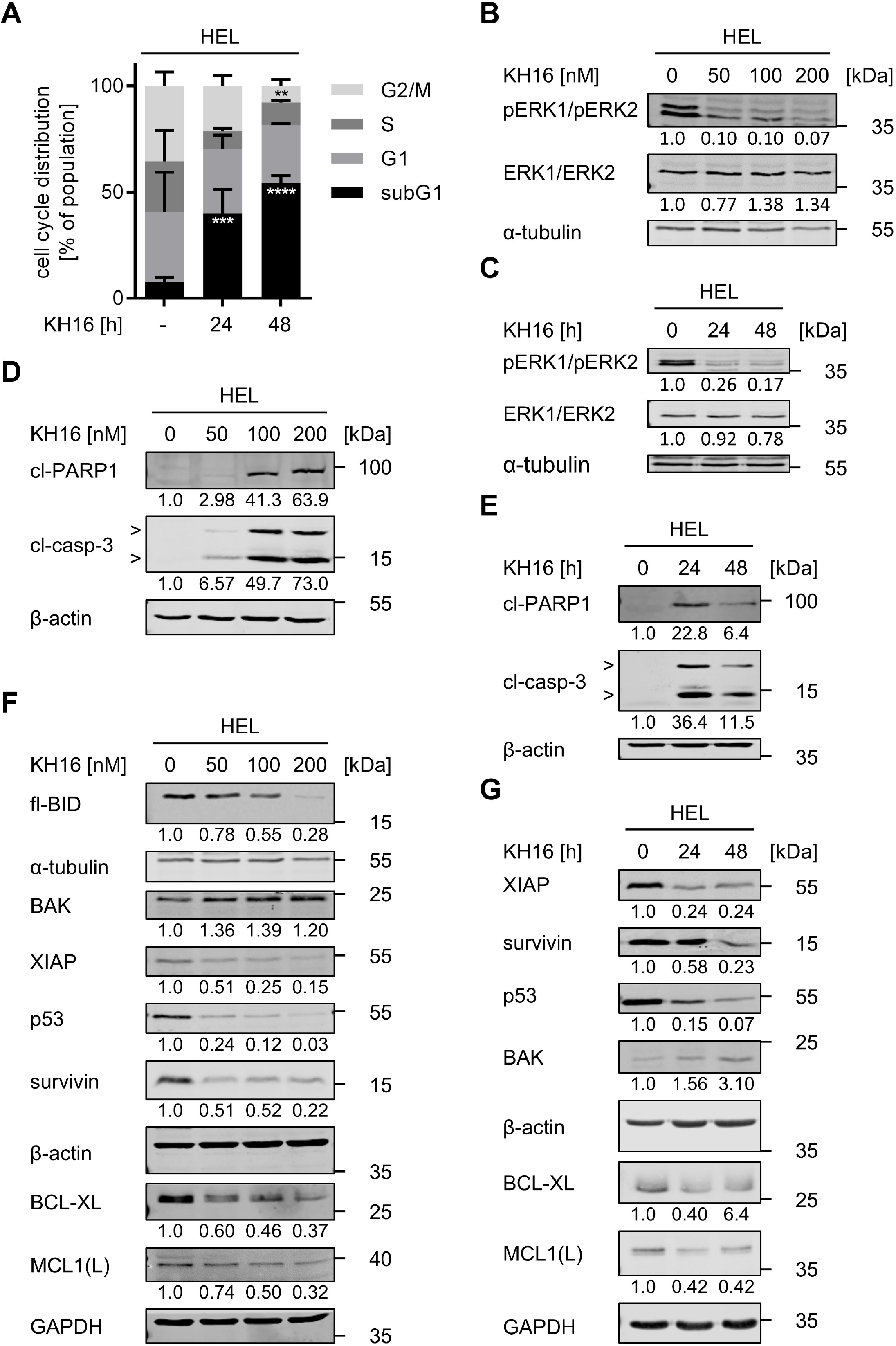
KH16 dysregulates the cell cycle and apoptosis regulators in HEL cells. (A) HEL cells were treated with 100 nM KH16 for 24 h or 48 h; -, untreated cells. Cells were fixed, stained with PI, and analyzed by flow cytometry. Results are derived from three independent experiments. Statistical tests for cell cycle analysis were performed using two-way ANOVA with Bonferroni correction (* p<0.05, ** p<0.01, *** p<0.001, **** p<0.0001). (B) KH16 blocks ERK phosphorylation. HEL cells were treated with 50, 100, or 200 nM KH16 for 24 h. Immunoblots were performed with antibodies against ERK1/ERK2, phosphorylated ERK1/ phosphorylated ERK2, and α-tubulin as loading control. (C) same as in B) but with 100 nM KH16 applied for 24-48 h. (D) KH16 activates caspase-3 and cleavage of its substrate PARP1 concentration-dependently. HEL cells were treated with 50, 100, or 200 nM KH16 for 24 h. Immunoblots were performed with antibodies against cleaved caspase-3, cleaved PARP1, and β-actin as loading control; cl, cleaved; casp, caspase. (E) Time-dependent activation of caspase-3 and PARP1 processing by 100 nM KH16. HEL cells were treated with 100 nM KH16 for 24 h or 48 h. Immunoblots were performed with antibodies against cleaved caspase-3, cleaved PARP1, and β-actin as loading control. (F,G) Dose- and time-dependent dysregulation of apoptosis regulators by KH16. HEL cells were treated with 50, 100, or 200 nM KH16 for 24 h (F), or 100 nM KH16 for 24 h to 48 h (G). Immunoblots were performed with antibodies against BID, BAK, XIAP, p53, survivin, BCL-XL, MCL1, with α-tubulin, β-actin and GAPDH as loading control; fl. full; MCL1(L), MCL1 large isoform. Quantifications were normalized to the loading control, untreated cells defined as 1.0; data are derived from two or more independent repetitions.

HEL cells have a constitutive activity of mitogen-activated protein kinases and extracellular signal-regulated kinases (MAPK/ERK1/ERK2). These promote cell cycle entry and unrestricted cell proliferation [35]. Treatment with 50 nM KH16 for 24 h strongly decreased the phosphorylation of ERK1/ERK2 at T202/Y204. This was not further augmented by 100-200 nM KH16. Total ERK1/ERK2 levels remained unchanged by KH16 (**Fig. 3B**). 100 nM diminished pERK1/pERK2 at 24 h and this effect persisted up to 48 h (**Fig. 3C**).

These results show that nanomolar concentrations of KH16 stall cell cycle proliferation, cause cytotoxic DNA damage, and suppress proliferative signaling through ERK1/ERK2.

### KH16 dysregulates apoptosis regulators

Apoptosis is executed upon the limited proteolysis of pro-caspases into active enzymes. Such activated caspase-3 is the ultimate death executioner at a “point-of-no-return” [15, 36]. This holds for HDACi-induced apoptosis [1, 4]. We treated HEL cells with 50-200 nM KH16 for 24 h and 100 nM KH16 for 24 h and 48 h and probed lysates from these cells for cleaved caspase-3 by immunoblot. We found a dose-dependent accumulation of active, cleaved caspase-3 fragments. Consistent herewith, we noted an accumulation of the cleaved form of the DNA repair protein PARP1 (**Fig. 3D**), which is a typical target of active caspase-3 in HDACi-treated tumor cells [32]. We further noted a time-dependent activation of caspase-3 and subsequent cleavage of PARP1 (**Fig. 3E**).

Next, we investigated the underlying mechanisms of KH16-induced apoptosis in HEL cells by immunoblot. Pro-apoptotic and anti-apoptotic proteins of the BCL2-family control mitochondrial integrity and thereby mitochondrially mediated apoptosis [15, 36]. An application of 50-200 nM KH16 for 24 h dose-dependently and significantly attenuated the levels of the anti-apoptotic BCL2 family members BCL-XL and MCL1 (**Fig. 3F**). Full-length BID was processed, which propels apoptosis. KH16 additionally reduced the anti-apoptotic proteins XIAP and survivin. An application of 50 nM KH16 for 24 h attenuated XIAP and survivin as effective as 100-200 nM KH16 (**Fig. 3F**). These effects persisted and increased upon a 48-h treatment period (**Fig. 3G**). The pro-apoptotic BAK is activated by truncated BID (tBID) and forms pores in mitochondrial membranes to trigger release of cytochrome-c for caspase activation [37]. BAK was expressed in resting HEL cells and KH16 augmented BAK levels after a treatment for 48 h (**Fig. 3G**).

The transcription factor p53 suppresses aberrant cell growth, but mutants thereof promote tumorigenesis and therapy resistance [38–40]. HEL cells carry p53 with an inactivating M133K mutation in its DNA-binding domain [41, 42]. KH16 reduced this mutant p53 protein dose- and time-dependently (**Fig. 3F,3G**).

These results illustrate that KH16 decreases anti-apoptotic proteins and induces the pro-apoptotic BAK. This ties in with caspase-dependent apoptosis.

### HDAC3 is a valid target in blood cancer cells

Of the new HDACi of the KH-series, KH16 is the most effective inhibitor of HDAC3 and the most effective inducer of apoptosis (**Table 1**, **Fig. 1**). We therefore analyzed whether HDAC3 is a dependency factor in various leukemic cells using data from the DepMap project. This database includes genome-wide CRISPR-Cas9 screens across a large panel of well characterized human tumor cell lines (https://depmap.org/portal/). The dependency of a cell line on a gene is reported as a dependency score, which represents the effect of the gene knockout on cell growth and viability. A score of zero indicates that a cell line is not dependent on the gene, whereas (strongly) negative scores indicate that the gene is essential in the given cell line. Analysis of the DepMap Crispr-Cas9 dataset identified that both myeloid and lymphoid tumor cells depend on HDAC3 for their proliferation and/or survival (**Fig. 4A**). The transcription factor p53 is a major tumor suppressor and mutated in 5-40% of aggressive myeloid and lymphoid cancer subtypes [38–40]. Like p53 wild-type leukemic cells, p53 mutant cells depend on HDAC3 (**Fig. 4A**).

**Figure 4:**
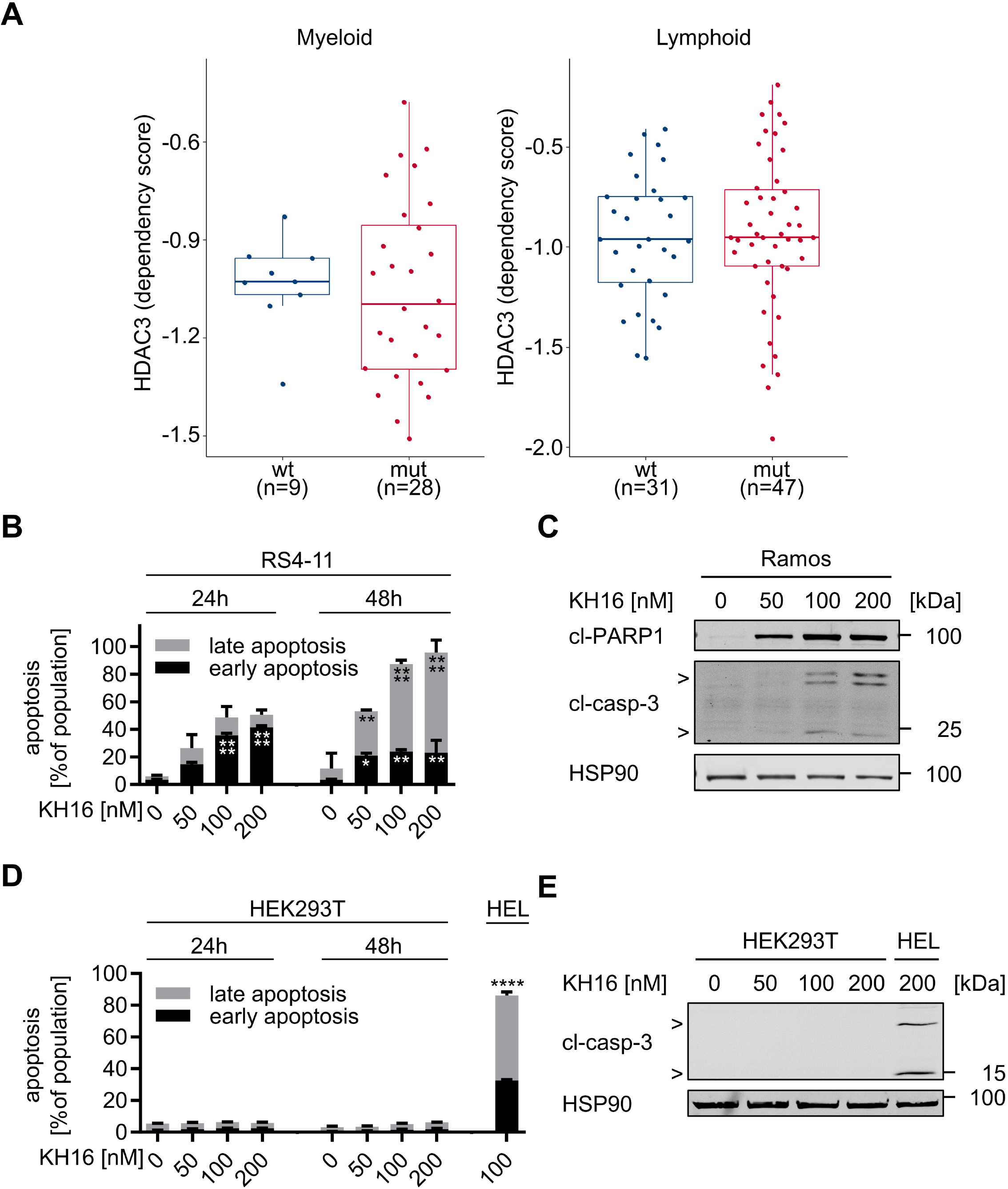
Myeloid and lymphoid leukemic cells rely on HDAC3 and are susceptible to KH16. (A) DepMap data analysis of HDAC3 dependencies in a panel of myeloid and lymphoid cell lines. Cell lines were stratified according to their TP53 status into p53 wild-type and p53 mutant cell lines. The boxplot shows the interquartile range in the box with the median as a horizontal line, the whiskers extend to 1.5 times the interquartile range. Each dot represents one cell line. p-values were determined using the Wilcoxon rank-sum test. A dependency score of 0 indicates that HDAC3 is not essential in each cell line, whereas a dependency score < 0 indicates that the cell line is more likely to be dependent on HDAC3. The median score of all common essential genes is -1. The Wilcoxon test was used to assess if there is a statistically significant difference in the HDAC3 dependency between p53 wild-type and p53 mutant cell lines. The p53 status has no statistically significant impact on the HDAC3 dependency: p=0.77 for myeloid and p=0.66 for lymphoid tumor cells. (B) RS4-11 cells were treated with 50, 100, or 200 nM KH16 for 24 or 48 h; n=3. Statistical tests were performed using two-way ANOVA test with Bonferroni correction (* p<0.05, ** p<0.01, **** p<0.0001). (C) Ramos cells were treated with 50, 100, or 200 nM KH16 for 24 h. Immunoblots were performed with antibodies against cleaved caspase-3 and cleaved PARP1 with HSP90 as loading control; cl, cleaved; casp, caspase. (D) The impact of 50, 100, and 200 nM KH16 on HEK293T cells after 24 h and 48 h was determined by annexin-V/PI-staining and flow cytometry (n=3). HEL cells treated with 100 nM KH16 for 48 h were used as positive control for apoptosis induction (n=2); 2-way ANOVA, Bonferroni correction; **** p<0.0001. (E) HEK293T and HEL cells were treated with 50, 100, or 200 nM KH16 for 24 h. Immunoblots were performed with antibodies against cleaved caspase-3 and cleaved PARP1, and HSP90 as loading control. Experiments represent two or more independent repetitions.

Due to these findings and to extend our data for the efficacy of KH16 against HEL cells, we assessed whether it induces apoptosis in human RS4-11 acute lymphoblastic leukemia cells (p53 wild-type) and Ramos B-cell lymphoma cells (p53 mutant) [43, 44]. KH16 activated caspase-3, the processing of PARP1, and annexin-V/PI-positivity in these cells (**Fig. 4B,4C; Fig. S13**) indicating efficient apoptosis induction.

To exclude that KH16 has general and non-selective killing effects, we incubated human embryonic kidney cells with KH16. These cells did not become apoptotic after 48 h when KH16 doses up to 200 nM KH16 were applied to them (**Fig. 4D**). Consistent herewith, no activation of caspase-3 in response to KH16 was observed in these cells (**Fig. 4E**). These data show that HDAC3 is a rational drug target in leukemic cells.

### KH16 suppresses autophagy protein expression

Autophagy is initiated by the formation of autophagosomes, which require catalytically active Unc-51 like autophagy activating kinase (ULK1), activated LC3B, and beclin-1. The sequestosome 1 (p62/SQSTM1) protein is degraded inside the autophagosome [45]. This multifunctional protein sorts ubiquitinylated proteins together with itself for degradation by autolysosomes [16]. We found that KH16 dose-dependently reduced the autophagy substrate p62 after 24 h in HEL cells (**Fig. S14**). This effect was pronounced upon an incubation for 48 h (**Fig. 5A**). Moreover, there was a time-dependent reduction of the autophagy-inhibiting phosphorylation of ULK1 at S757 and a decrease of the autophagy-promoting protein beclin-1 by 100 nM KH16 (**Fig. 5A**). Activation of LC3B manifests as phosphatidylethanolamine-conjugated band that migrates faster in immunoblot analyses. This conversion of LC3-I to LC3-II denotes autophagosome formation [45]. We did not see activation of LC3B (**Fig. 5A**). Since the reduction of the inhibitory phosphorylation of ULK1 may still indicate autophagy, we suppressed it with chloroquine. This compound prevents the autophagosome-lysosome fusion and thereby interrupts the autophagic flux [46]. Chloroquine did not alter the KH16-induced activation of caspase-3 and PARP1 cleavage (**Fig. 5B**).

**Figure 5:**
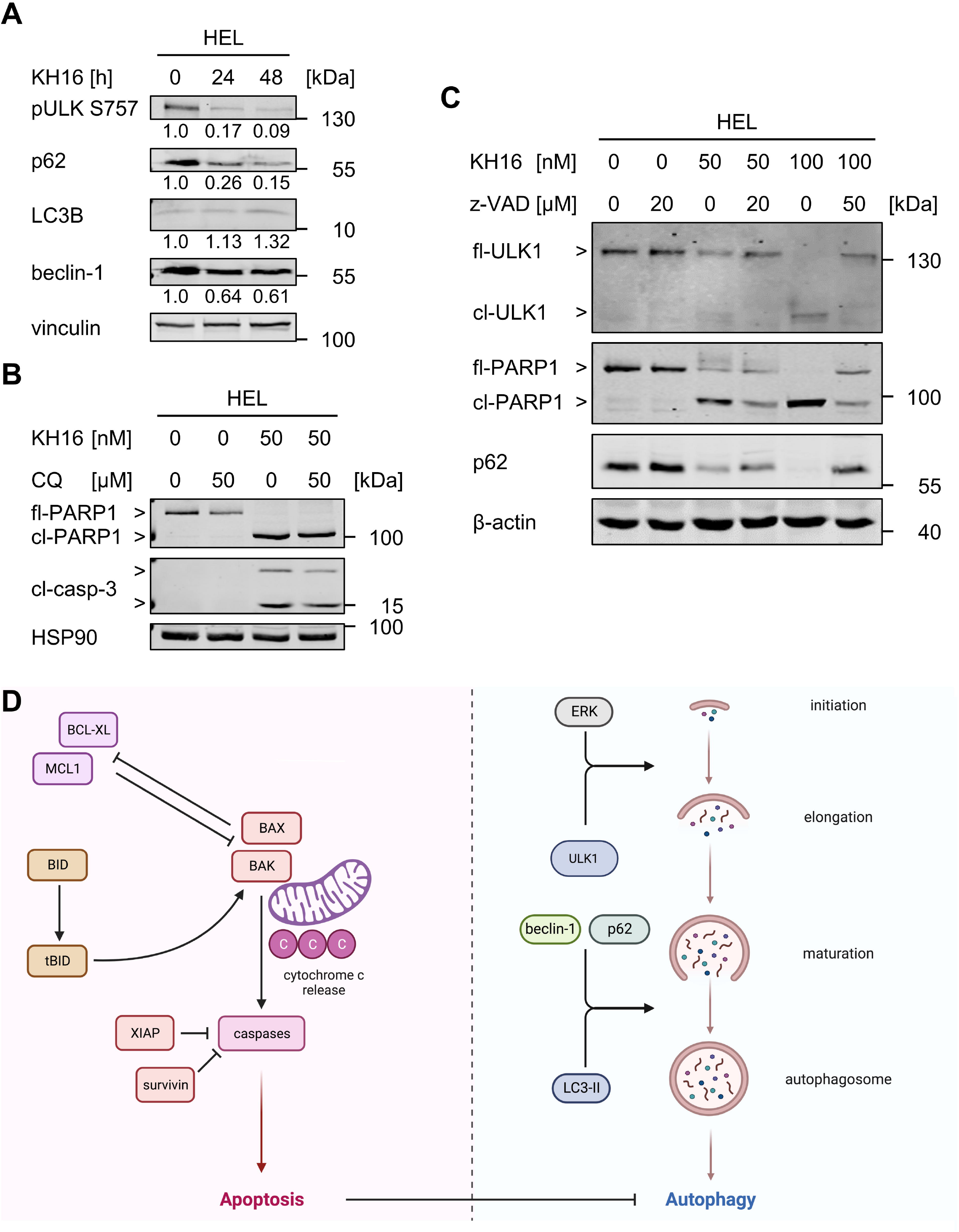
KH16 modulates autophagy and apoptosis in leukemic cells. (A) HEL cells were treated with 100 nM KH16 for 24 h to 48 h. Immunoblots were performed with antibodies against phosphorylated ULK1, p62, LC3B, beclin-1, and vinculin as loading control. (B) HEL cells were treated with 50 nM KH16 and/or 50 µM chloroquine for 24 h. Immunoblots were performed against cleaved caspase-3, PARP1, and HSP90 as loading control; CQ, chloroquine; fl, full length; cl, cleaved; casp., caspase. (C) HEL cells were treated with 50 or 100 nM KH16, and/or 20 or 50 µM z-VAD-FMK for 24 h. Immunoblots were performed with antibodies against ULK1 serine 757, p62 and PARP1; β-actin as loading control. Quantifications were normalized to the loading control, untreated cells defined as 1.0. Data represent the outcomes of two or more independent repetitions. (D) Overview of the KH16-induced cell death. A key step promoted by cleavage of BID into pro-apoptotic tBID is the formation of the BAX/BAK oligomer protein. Oligomerization of BAX and BAK facilitates the release of cytochrome-c from the mitochondrial membrane. BCL-XL and MCL-1 directly inhibit the BAX/BAK oligomer formation at the mitochondrial membrane. The anti-apoptotic proteins XIAP and survivin suppress the activation of caspase-3. Release of cytochrome-c from the mitochondrial membrane activates caspases suppressing autophagy.

These data let us speculate that the reduction of p62 and p-ULK1 by KH16 is not indicative of increased autophagy. Since KH16 causes apoptosis (**Figs. 1,3**), we speculated that caspase activation decreased p62 and ULK1. Like the established caspase-3 target PARP1, ULK1 was processed into a smaller protein fragment in KH16-treated HEL cells. This effect occurred dose-dependently. The pan-caspase inhibitor z-VAD-FMK restored full-length ULK1 and prevented its processing into the smaller fragment. Z-VAD-FMK also restored the levels of p62 in such cells (**Fig. 5C**). **Fig. 5D** summarizes the key findings of this study on the mechanisms through which KH16 triggers apoptosis in leukemic cells. We unravel that major autophagy-inducing proteins are cleaved by caspases in KH16-treated cells. From these data, we conclude that KH16 has a negative impact on autophagy through activation of caspases and a subsequent degradation of autophagy proteins.

## Discussion

The FDA has approved HDACi as treatment option for cutaneous T cell lymphoma and myeloma [1,4,5]. MPNs mainly comprise polycythemia vera (PV), essential thrombocythemia (ET), primary myelofibrosis (PMF), and chronic myeloid leukemia (CML) [2]. Median survival rates in patients affected by MPNs range from 19.8 years in patients with ET to 13.5 years in patients diagnosed with PV and 5.9 years in patients with PMF. The transition of such diseases to acute myeloid leukemia (AML) and thrombotic events are often fatal. Thus, additional treatment options to the established options hydroxyurea and interferons are required to control MPNs. The hydroxamic acid-based HDACi panobinostat and givinostat produced promising results in MPN patients [47, 48]. KH16 might be an additional HDACi in the treatment regimen of MPNs. Since KH16 is effective against human acute lymphoblastic leukemia and B-cell lymphoma cells from difficult-to-treat leukemia [43, 44], it may also be a promising drug for treating such diseases. Remarkably, KH16 has no negative impact on human embryonic kidney cells, suggesting a specific anti-tumor cell effect.

In vitro and in cells, HDACi of the KH series are more potent inducers of apoptosis than their parental structure SAHA [6]. KH16 is the most potent of them and has the strongest activity against HDAC3. HDAC3 has key survival functions in leukemic cells of various hematopoietic lineages. These include the control of cell proliferation, apoptosis, DNA replication fork stability, and genomic integrity [31,49–55]. Intriguingly, a full-body knockout of HDAC3 was not lethal in mice [56]. These data together with CRISPR-Cas9 screening data from a panel of myeloid and lymphoid cancer cell lines suggest that HDAC3 is a particular vulnerability of blood cancer cells and that future research on HDACi should focus on agents with strong activity against HDAC3. The narrower inhibition profile of KH16 compared to established HDACi, combined with its high anti-proliferative activity suggest further testing of KH16 and derivatives thereof. According to our results, these should contain an unmethylated indole in the capping group for interaction with HDAC3.

Previous studies showed that class I HDAC inhibition reduced p53 levels [32,57–60]. Congruently, KH16 attenuates p53 levels. Wild-type p53 acts as a tumor suppressor gene with the capability to induce cell death via apoptosis. HEL cells carry the p53^M133K^ mutation rendering p53 inactive [41]. It remains to be shown whether this p53 mutant induces survivin and BCL-XL like p53^R172H^ in murine tumor cells (corresponds to the human p53 hotspot mutation p53^R175H^) [58, 59]. If this applies, reduction of p53^M133K^ might have caused the attenuation of survivin and BCL-XL in HEL cells. Irrespective thereof, our analyses demonstrate that leukemic cells rely on HDAC3. Hence, patients with different p53 status may profit from KH16 and other HDACi with a strong inhibitory impact on HDAC3.

KH16 mainly reduced cells in the S- and G2/M-phases and had a lesser impact on cells in G1 phase. The stronger sensitivity of such cells is consistent with the literature and multiple explanations for this have been provided for leukemic cells. These include, but are not limited to, an inhibition of the S phase-dependently induced transcription factor NF-κB [59,61,62], the induction of DNA replication stress/DNA damage [31,49–51,53], inactivation of signaling through the transcription factor STAT5 [63], and a disruption of proper mitosis [64]. KH16-induced cell cycle disruptions are linked to a significant fragmentation of DNA. Activated ERK1/ERK2 regulate gene expression driving cell cycle progression as well as resistance towards kinase inhibitors [35, 65]. Albeit we did not specifically investigate other proteins in the MAPK/ERK pathway, our observations suggest the inactivation of ERKs by KH16 is a mechanism contributing to cell cycle arrest and apoptosis.

Apoptosis appears as the main anti-leukemic mechanism that is induced by KH16. In some experiments, we noticed cleavage of PARP1 before a detectable activation of caspase-3. This can be explained by a processing of PARP1 by additional apoptosis-promoting enzymes, such as caspase-7 [66, 67]. In addition to apoptosis, HDACi modulate autophagy in leukemic cells [5]. Opposed to the destruction of cells by apoptosis, autophagy is characterized by self-digestion through autophagosomes. This frequently protects cells from HDACi-induced apoptosis [5,17–19,45]. KH16 decreases pULK1 Ser757 which suppresses autophagy as well as the autophagy inducers p62 and beclin-1. An activation of ULK1 and attenuation of p62 can indicate ongoing apoptosis [16, 45]. However, experiments with chloroquine suggest that autophagy plays a minor role in the KH16-mediated cell death. Instead, KH16 induces the caspase-mediated processing of ULK1 and p62. Thus, apoptosis can operate as an upstream regulator of autophagy in HDACi-treated leukemic cells.

The induction of autophagy can be an undesired effect of HDACi. For example, SAHA induces autophagy in CML cells. This process restricts HDACi-induced apoptosis that occurs without functional p53 but dependent on the apoptosis-promoting lysosomal protease cathepsin-D [68]. Unlike in CML cells, the fatty acid-based class I HDACi valproic acid, the benzamide entinostat, and SAHA suppress autophagy in pediatric AML cells [69]. Both studies found that HDACi induced an accumulation of reactive oxygen species and DNA damage. Also, in pre-B acute lymphocytic leukemia-derived cells autophagy attenuates pro-apoptotic effects of the pan-HDACi panobinostat [70]. The HDAC1/2/3/11 inhibitor mocetinostat and the HDAC1/2/3/10 inhibitor tucidinostat kill chronic lymphocytic leukemia B-cells through apoptosis and a suppression of autophagy [71, 72]. Albeit HDAC10 regulates autophagy in leukemic cells, cytotoxic effects due to HDAC10 inhibition unlikely explain our data. At least in AML cells, an induction of autophagy upon a specific inhibition of HDAC10 does not cause apoptosis [73]. It appears that KH16 is superior to HDACi that propel cytoprotective autophagy. These findings suggest further investigation of KH16 as a drug for the treatment of blood cancers and potentially other tumor types.

## Conclusions

HDACi of the KH-series kill leukemic cells, outperforming the clinically validated HDACi SAHA in preclinical experiments. Defined chemical structures can deliver a pharmacology to preferentially inhibit the cancer-relevant HDAC3. There is a previously unrecognized acetylation-regulated, caspase-dependent upstream control of autophagy by apoptosis. KH16 triggers apoptosis and caspase activity that eliminates autophagy proteins.

## Figure legends

**S13:**
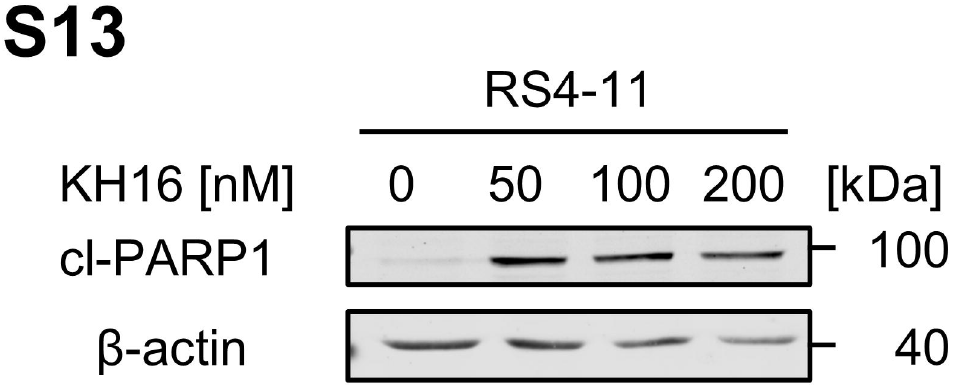
Apoptotic effect of KH16 on RS4-11 cells. RS4-11 were treated with 0, 50, 100, and 200 nM KH16 for 24 h. Immunoblots with antibodies against cleaved PARP1 (cl, cleaved) and β-actin as loading control, experiment represents two repetitions; cl, cleaved.

**S14:**
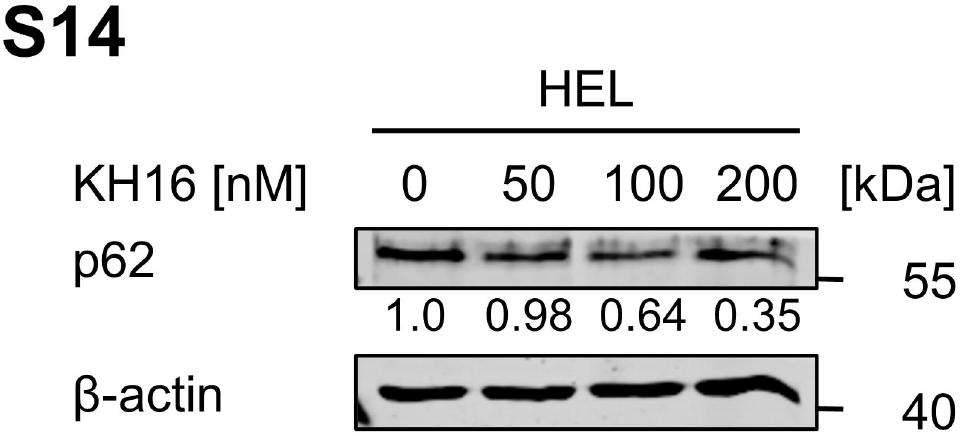
Effect of KH16 on p62 in HEL cells. HEL cells were treated with 0, 50, 100, and 200 nM KH16 for 24 h. Expression of p62 was analyzed by immunoblot; β-actin as loading control. Quantifications were normalized to the loading control, untreated cells defined as 1.0, experiment represents two independent repetitions.

## Author contributions

M.A.F. performed most experiments. A.M.M., R.A., A.G.K., M.C.L., C.B., and A.P-S. did additional experiments. A.M.M. and O.H.K. contributed the scientific development, discussion, and supervision of the project. K.H. and H.I.S. synthesized and characterized the KH compounds and carried out the ligand docking. M.Z. and M.S. carried out the enzymatic testing. M.C.L. analyzed the DepMap datasets. T.G.H. secured funding of M.C.L. and commented the manuscript. W.S. designed the synthesis of KH compounds for biological experiments and wrote the parts on chemistry in the manuscript. M.A.F. drafted the manuscript and O.H.K. finalized the manuscript with the help of W.S.

## Note

This work contains substantial parts of the Bachelor thesis of M.A.F.

## Abbreviations

AML: acute myeloid leukemia
BAK: BCL2 homologous antagonist/killer
BAX: BCL2-like protein-4
BCL2: B-cell lymphoma-2
BCL-XL: B-cell lymphoma extra-large
BID: BH3 interacting-domain death agonist
BIM: Bcl-2-like protein-11
BSA: bovine serum albumin
CML: chronic myeloid leukemia
ClQ: chloroquine
cyt-c: cytochrome c
ERK: extracellular-signal regulated kinase
ET: essential thrombocythemia
FCS: fetal calf serum
FDA: food- and-drug-administration office
GAPDH: glyceraldehyde 3-phosphate dehydrogenase
HEPES: 4-(2-hydroxyethyl)-1-piperazineethanesulfonic acid
HDACs: histone deacetylases
HDACi: histone deacetylase inhibitor(s)
HSP90: heat shock protein 90kDa
MAPK: mitogen-activated protein kinase
MCL1: myeloid cell leukemia sequence-1 (BCL2-Related)
MPNs: myeloproliferative neoplasms
PARP1: poly-ADP-ribose polymerase 1
PI: propidium iodide
PMF: primary myelofibrosis
PV: polycythemia vera
ULK1: unc-51 like autophagy activating kinase
SAHA: suberoylanilide hydroxamic acid
STAT5: signal transducer and activator of transcription-5
TCEP: tris(2-carboxyethyl)phosphine
XIAP: X-linked inhibitor of apoptosis protein
ZMAL: benzyl {6-acetamido-1-[(4-methyl-2-oxo-2H-chromen-7-yl)amino]-1-oxohexan-2-yl}carbamate

## Acknowledgement

We thank Prof. Dr. Thorsten Heinzel, Prof. Dr. F.-D. Böhmer, Dr. Kosan, Jena, and Dr. Manuel Grez, Frankfurt/Main for cell lines.

## Declarations

### Availability of data and material

Data generated or analyzed during this study and material are available from the corresponding authors upon reasonable scientific request.

---

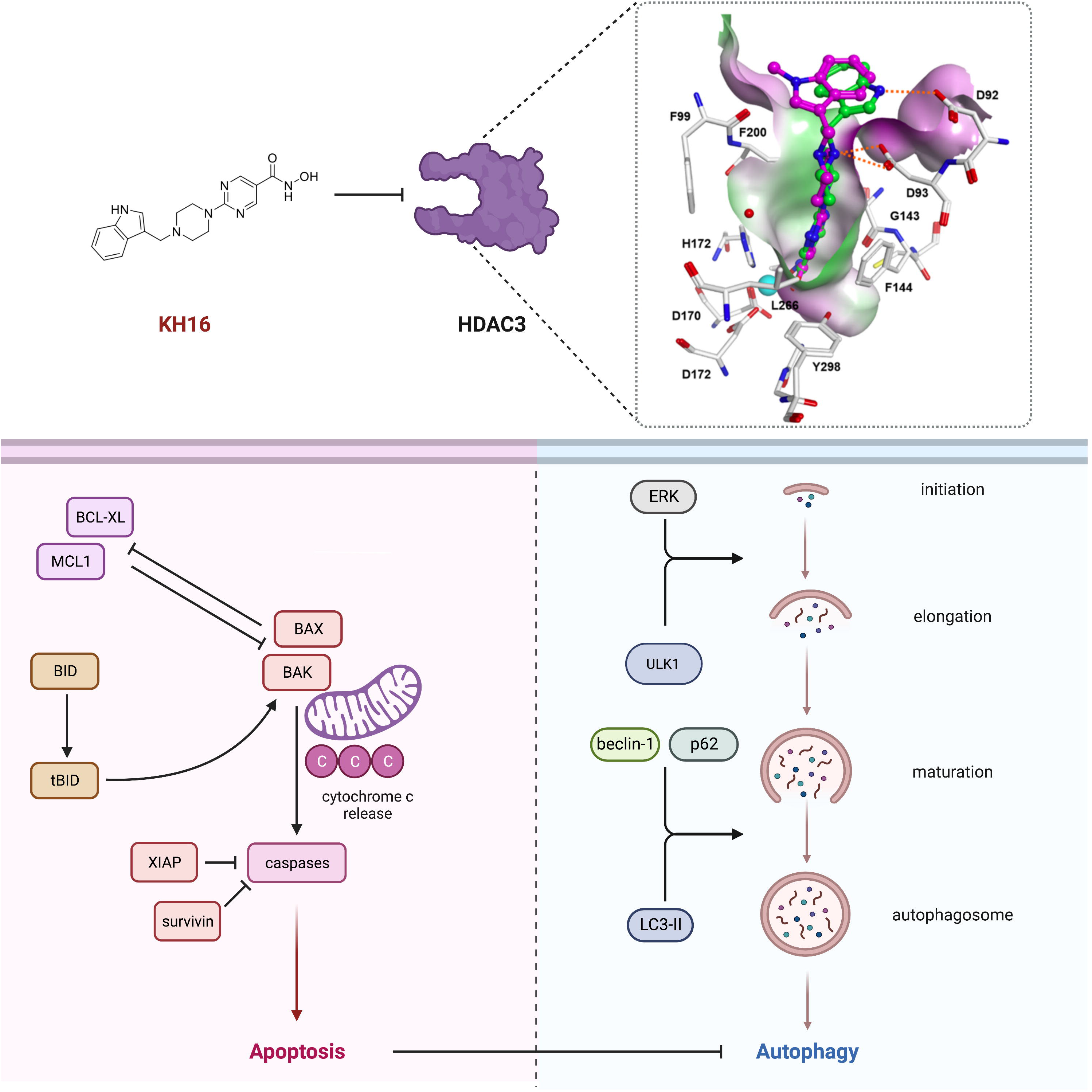

## Chemical synthesis

The synthesis of the compounds KH9 (**7a),** KH16 **(7b) and** KH29 **(7c)** is depicted in Scheme 1. Initially ester **3** was obtained by nucleophilic aromatic substitution with commercially available ethyl 2-chloropyrimidine-5-carboxylate **1** and tert-butyl piperazine-1-carboxylate **2**. Subsequent deprotection of the tert-butoxycarbonyl protecting group led to amine **4**. Reductive amination with the respective aldehydes gave intermediates **5a-c** which were hydrolysed under alkaline conditions to yield carboxylic acids **6a-c**. Coupling of O-(tetrahydro-2H-pyran-2-yl)hydroxylamine and following deprotection of the 2-tetrahydropyranyl group under acidic conditions gave the final hydroxamic acids **7a-c**.

**Scheme 1.**
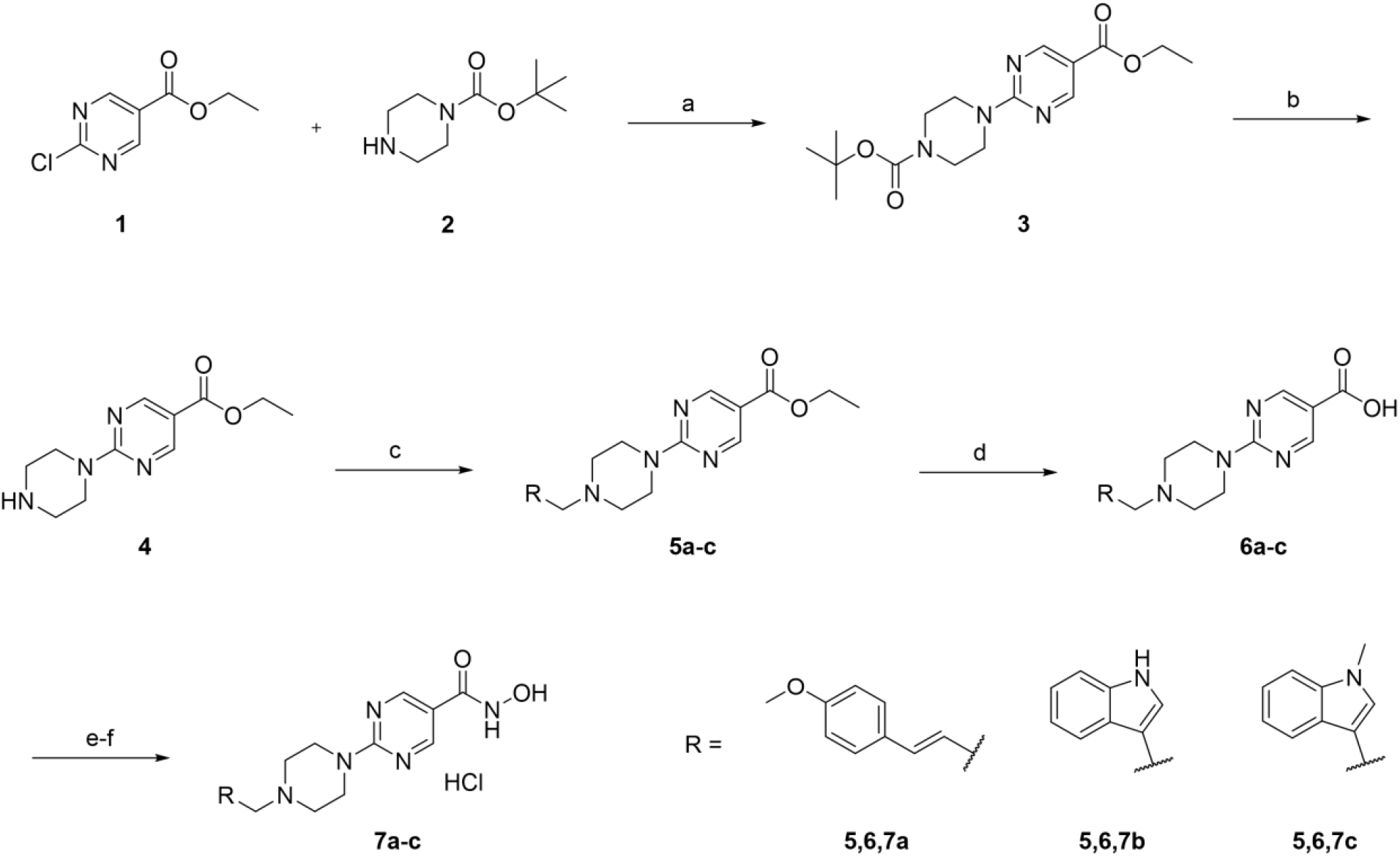
Synthesis of the compounds 6a-c. Reagents and conditions: a) DIPEA, CH_2_Cl_2_, rt, 3h; b) TFA/CH_2_Cl_2_, rt, 30 min; c) RCHO, NaBH(OAc)_3_, AcOH, CH_2_Cl_2_, rt, overnight ; d) i) 1M NaOH, MeOH, reflux, 1h, ii) 1M HCl, 0°C, 30 min; e) H_2_NOTHP, PyBOP, DIPEA, THF, rt, overnight; f) HCl, MeOH, rt, overnight

### Synthesis of Ethyl 2-(4-(tert-butoxycarbonyl)piperazin-1-yl)pyrimidine-5-carboxylate **(3)**

To a solution of tert-butyl piperazine-1-carboxylate (1996 mg, 10.718 mmol, 1 eq.) and ethyl-2-chloropyrimidine-5-carboxylate (2000 mg, 10.718 mmol,1 eq.) in CH_2_Cl_2_ was added DIPEA (4.67 mL, 26.795 mmol, 2.5 eq.) and the reaction mixture was stirred at room temperature for 3h. The reaction mixture was washed with water and the organic phase was dried over anhydrous sodium sulfate, filtered, and evaporated in vacuo to dryness. The solid crude product **3** was directly used without further purification. 3523 mg (10.472 mmol; 98 %) ^1^H NMR (400 MHz, CDCl_3_): δ 8.84 (s, 2H), 4.34 (q, J = 7.1 Hz, 2H), 3.99 – 3.82 (m, 4H), 3.60 – 3.42 (m, 4H), 1.49 (s, 9H), 1.36 (t, J = 7.1 Hz, 3H). ESI-MS m/z: 359.32 [M+Na]^+^.

### Synthesis of Ethyl 2-(piperazin-1-yl)pyrimidine-5-carboxylate **(4)**

Compound **3** was dissolved in a mixture of TFA and CH_2_Cl_2_ (1:2) and stirred at room temperature for 30 min. After the solvent was removed from the reaction mixture in vacuo, the residue was taken up in CH_2_Cl_2_ and washed with 1M aqueous sodium bicarbonate solution. The organic phase was dried over anhydrous sodium sulfate, filtered and the solvent was evaporated in vacuo to dryness The crude product **4** was used in the next reaction step without further purification. 2400 mg (10.155 mmol; 97 %) ^1^H NMR (400 MHz, d6-DMSO): δ 8.75 (s, 2H), 4.26 (q, J = 7.1 Hz, 2H), 3.78 (dd, J = 5.9, 4.4 Hz, 4H), 3.25 (s, 1H), 2.76 (dd, J = 5.9, 4.3 Hz, 4H), 1.28 (t, J = 7.1 Hz, 3H). ESI-MS m/z: 237.64 [M+H]^+^.

### General procedure for the synthesis of compounds 5a-c

To a solution of compound **4** (400 mg, 1.693 mmol, 1 eq.) in CH_2_Cl_2_ was added the respective aldehyde (1.693 mmol; 1 eq.) and glacial acetic acid (97µL, 1.693 mmol, 1 eq.). The reaction mixture was stirred for 15 min at room temperature. After addition of sodium triacetoxyborohydride (539 mg, 2.540 mmol, 1.5 eq.) the reaction mixture was stirred overnight at room temperature. The reaction mixture was then poured into saturated aqueous sodium bicarbonate solution and stirred for 15 min at room temperature. The organic layer was separated, dried over anhydrous sodium sulfate, filtered and the solvent was evaporated in vacuo to dryness. The crude product obtained was purified by flash column chromatography (SiO_2_, EA/PE).

### Ethyl (E)-2-(4-(3-(4-methoxyphenyl)allyl)piperazin-1-yl)pyrimidine-5-carboxylate (**5a**)

Compound **5a** was synthesized according to the above general procedure. 523 mg (1.368 mmol; 81%); ^1^H NMR (CDCl_3_, 400 MHz): δ 8.83 (s, 2H), 7.38 – 7.29 (m, 2H), 6.91 – 6.80 (m, 2H), 6.48 (d, J = 15.8 Hz, 1H), 6.14 (dt, J = 15.8, 6.8 Hz, 1H), 4.33 (q, J = 7.1 Hz, 2H), 4.07 – 3.91 (m, 4H), 3.81 (s, 3H), 3.18 (d, J = 6.8 Hz, 2H), 2.64 – 2.50 (m, 4H), 1.36 (t, J = 7.1 Hz, 3H). ESI-MS m/z: 383.76 [M+H]^+^.

### Ethyl 2-(4-((1H-indol-3-yl)methyl)piperazin-1-yl)pyrimidine-5-carboxylate (**5b**)

Compound **5b** was synthesized according to the above general procedure. 518 mg (1.417 mmol; 84 %); ^1^H NMR (CDCl_3_, 400 MHz): δ 8.81 (s, 2H), 8.12 (s, 1H), 7.83 – 7.69 (m, 1H), 7.45 – 7.30 (m, 1H), 7.25 – 7.19 (m, 1H), 7.18 – 7.10 (m, 2H), 4.33 (q, J = 7.1 Hz, 2H), 4.03 –3.89 (m, 4H), 3.78 (s, 2H), 2.68 – 2.49 (m, 4H), 1.36 (t, J = 7.1 Hz, 3H). ESI-MS m/z: 366.40 [M+H]^+^.

### Ethyl 2-(4-((1-methyl-1H-indol-3-yl)methyl)piperazin-1-yl)pyrimidine-5-carboxylate (**5c**)

Compound **5c** was synthesized according to the above general procedure. 508 mg (1.339 mmol; 79 %) ^1^H NMR (CDCl_3_, 400 MHz): δ 8.81 (s, 2H), 7.79 – 7.67 (m, 1H), 7.37 – 7.29 (m, 1H), 7.25 – 7.21 (m, 1H), 7.16 – 7.10 (m, 1H), 7.02 (s, 1H), 4.33 (q, J = 7.1 Hz, 2H), 3.94 (s, 4H), 3.78 (s, 3H), 3.77 (s, 2H), 2.58 (s, 4H), 1.36 (t, J = 7.1 Hz, 3H). ESI-MS m/z: 380.04 [M+H]^+^.

### General procedure for the synthesis of compounds **6a-c**

The carboxylic acid ester **5a-c** (1 eq.) was dissolved with MeOH and 1M aqueous sodium hydroxide solution (2.5 eq.). The reaction mixture was refluxed for 1h. Then the solvent was removed under reduced pressure and the residue was taken up in water and acidified with 1M aqueous hydrochloric acid at 0°C until the carboxylic acid was obtained as precipitate. The precipitate was isolated by filtration, washed with water and dried. The crude product **6a-c** was used without further purification.

### (E)-2-(4-(3-(4-methoxyphenyl)allyl)piperazin-1-yl)pyrimidine-5-carboxylic acid (**6a**)

Compound **6a** was synthesized from **5a** (409 mg, 1.068 mmol) according to the above general procedure. (347 mg, 0.979 mmol, 92%) ^1^H NMR (DMSO-d_6_, 400 MHz): δ 12.98 (s, 1H), 8.83 (s, 2H), 7.60 – 7.25 (m, 2H), 7.05 – 6.87 (m, 2H), 6.82 – 6.54 (m, 1H), 6.37 – 5.98 (m, 1H), 4.79 (s, 2H), 3.82 (s, 2H), 3.77 (s, 3H), 3.66 – 2.75 (m, 6H). ESI-MS m/z: 353,51 [M-H]^-^ 2-(4-((1H-Indol-3-yl)methyl)piperazin-1-yl)pyrimidine-5-carboxylic acid (**6b**) Compound **6b** was synthesized from **5b** (518 mg, 1.417 mmol) according to the above general procedure. (415 mg, 1.230 mmol, 87%) ^1^H NMR (DMSO-d6, 400 MHz): δ.88 (s, 1H), 11.47 (s, 1H), 8.80 (s, 2H), 7.77 (d, J = 7.8 Hz, 1H), 7.60 (s, 1H), 7.42 (d, J = 8.0 Hz, 1H), 7.18 – 7.10 (m, J = 7.5 Hz, 1H), 7.10 – 7.03 (m, J = 7.4 Hz, 1H), 4.41 (s, 2H), 3.59 – 3.05 (m, 8H). ESI-MS m/z: 338,30 [M+H]^+^.

### 2-(4-((1-Methyl-1H-indol-3-yl)methyl)piperazin-1-yl)pyrimidine-5-carboxylic acid (**6c**)

Compound **6c** was synthesized from **5c** (510 mg, 1.318 mmol) according to the above general procedure. (408 mg, 1.160 mmol, 88%) ^1^H NMR (DMSO-d_6_, 400 MHz): δ 8.74 (s, 2H), 7.66 (d, J = 7.8 Hz, 1H), 7.39 (d, J = 8.2 Hz, 1H), 7.24 (s, 1H), 7.18 – 7.10 (m, 1H), 7.07 – 6.98 (m, 1H), 3.91 – 3.79 (m, 4H), 3.75 (s, 3H), 3.68 (s, 2H), 2.49 – 2.43 (m, 4H). ESI-MS m/z: 350,35 [M-H]^+^.

### General procedure for the synthesis of compounds **7a-c**

The respective carboxylic acid **6a-c** (1 eq) was dissolved in dry THF. PyBOP (1.2eq) and DIPEA (2.5 eq.) were added and the reaction mixture was stirred at room temperature for 15 min. Then NH_2_OTHP (1.5eq.) was added and the reaction mixture was stirred at room temperature overnight. The solvent was evaporated in vacuo and the residue was diluted with 1M aqueous sodium bicarbonate solution and extracted with ethyl acetate. The combined organic layers were dried over anhydrous sodium sulfate, filtered and the solvent was evaporated in vacuo to dryness. The crude product obtained was purified by flash column chromatography (SiO_2_, CHCl_3_/MeOH). The purified product was dissolved in MeOH and concentrated hydrochlorid acid (2.5 eq.) was added. After stirring at room temperature overnight, the formed precipitate was filtered from the reaction mixture and washed with CH_2_Cl_2_. If necessary the product was purified by flash column chromatography (SiO_2_, CHCl_3_/MeOH). Final analysis of purity was done by HPLC. All compounds showed purity above 98.9%.

### (E)-N-Hydroxy-2-(4-(3-(4-methoxyphenyl)allyl)piperazin-1-yl)pyrimidine-5-carboxamide Hydrchloride KH9 (**7a**)

Compound **7a** was synthesized from **6a** (346 mg, 0.977 mmol) according to the above general procedure. (160 mg, 0.394 mmol, 40%) ^1^H NMR (DMSO-d_6_, 400 MHz): δ 11.05 (s, 1H), 8.99 (s, 1H), 8.67 (s, 2H), 7.50 – 7.24 (m, 2H), 6.99 – 6.76 (m, 2H), 6.48 (d, J = 15.9 Hz, 1H), 6.16 (dt, J = 15.9, 6.6 Hz, 1H), 3.85 – 3.77 (m, 4H), 3.75 (s, 3H), 3.11 (d, J = 6.5 Hz, 2H), 2.48 – 2.42 (m, 4H). ^13^C NMR (DMSO-d_6_, 101 MHz): δ 161.37, 158.73, 157.02, 131.84, 129.24, 127.44, 124.21, 114.55, 113.96, 60.20, 55.07, 52.30, 43.45; HRMS (ESI): m/z calcd. for C_19_H_24_N_5_O_3_, [M+H]^+^ 370.1879, found 370.1872; HPLC: rt = 10.280 min. (Purity 99.54%).

### 2-(4-((1H-Indol-3-yl)methyl)piperazin-1-yl)-N-hydroxypyrimidine-5-carboxamide Hydrochloride KH16 (**7b**)

Compound **7b** was synthesized from **6b** (415 mg, 1.230 mmol) according to the above general procedure. (287 mg, 0. 738 mmol, 60%) ^1^H NMR (DMSO-d_6_, 400 MHz): δ 11.05 (s, 1H), 10.96 (s, 1H), 8.99 (s, 1H), 8.65 (s, 2H), 7.79 – 7.55 (m, 1H), 7.46 – 7.33 (m, 1H), 7.26 (s, 2H), 7.12 – 7.04 (m, 1H), 7.02 – 6.89 (m, 1H), 3.79 (s, 4H), 3.70 (s, 2H).). ^13^C NMR (DMSO-d6, 101 MHz): δ δ 161.85, 161.29, 157.03, 136.29, 127.54, 124.73, 120.97, 118.99, 118.48, 114.53, 111.35, 53.03, 52.10, 43.41. HRMS (ESI): m/z calcd. for C_18_H_21_N_6_O_2_ [M+H]^+^: 353.1726; found 353.172.; HPLC: rt = 4,851 min. (Purity 99.80 %).

### N-hydroxy-2-(4-((1-methyl-1H-indol-3-yl)methyl)piperazin-1-yl)pyrimidine-5-carboxamide Hydrochloride KH29 (**7c**)

Compound **7c** was synthesized from **6c** (400 mg, 1.138 mmol) according to the above general procedure. (237 mg, 0.588 mmol, 52%) ^1^H NMR (DMSO-d_6_, 400 MHz): δ 11.24 (s, 2H), 9.06 (s, 1H), 8.75 (s, 2H), 7.87 – 7.77 (m, 1H), 7.64 (s, 1H), 7.53 – 7.44 (m, 1H), 7.30 – 7.18 (m, 1H), 7.17 – 6.98 (m, 1H), 4.78 (d, J = 14.4 Hz, 2H), 4.47 (d, J = 4.4 Hz, 2H), 3.82 (s, 3H), 3.55 – 3.38 (m, 4H), 3.19 – 2.92 (m, 2H). ^13^C NMR (DMSO-d_6_, 101 MHz): δ 161.42, 161.02, 157.21, 136.38, 132.94, 127.86, 121.76, 119.79, 118.83, 115.73, 110.11, 101.38, 49.96, 49.40, 40.44, 32.71. HRMS (ESI): m/z calcd. for C_19_H_23_N_6_O_2_ [M+H]^+^: 367.1882; found 367,188; HPLC: rt = 8,387 min. (Purity 98.92%).

All materials and reagents were purchased from Sigma-Aldrich Co. Ltd. and abcr GmbH. All solvents were analytically pure and dried before use. Thin layer chromatography was carried out on aluminum sheets coated with silica gel 60 F254 (Merck, Darmstadt, Germany). For medium pressure chromatography (MPLC) silica gel 60 (0.036e0.200 mm) was used. Final compounds were confirmed to be of >95% purity based on HPLC. Purity was measured by UV absorbance at 254 nm. The HPLC consists of an XTerra RP18 column (3.5 µm, 3.9 mm x 100 mm) from the manufacturer Waters (Milford, MA, USA) and two LC-10AD pumps, a SPD-M10A VP PDA detector, and a SIL-HT autosampler, all from the manufacturer Shimadzu (Kyoto, Japan). Mass spectrometry analyses were performed with a Finnigan MAT710C (Thermo Separation Products, San Jose, CA, USA) for the ESIMS spectra and with a LTQ (linear ion trap) Orbitrap XL hybrid mass spectrometer (Thermo Fisher Scientific, Bremen, Germany) for the HRMS-ESI (high resolution mass spectrometry) spectra. For the HRMS analyses the signal for the isotopes with the highest prevalence was given and calculated (^35^Cl, ^79^Br). ^1^H NMR and ^13^C NMR spectra were taken on a Varian Inova 500 using deuterated chloroform or deuterated dimethylsulfoxide as solvent. Chemical shifts are referenced to the residual solvent signals. The following abbreviations and formulas for solvents and reagents were used: ethyl acetate (EtOAc), dimethylformamide (DMF), dimethylsulfoxide (DMSO), methanol (MeOH), tetrahydrofuran (THF), chloroform (CHCl_3_), water (H_2_O), dichloromethane (DCM), N,N-diisopropylethylamine (DIPEA), trimethylamine (TEA), hydrochloric acid (HCl) and trifluoroacetic acid (TFA).

## Analytical data

### KH9 (7a)

**Figure S 1:**
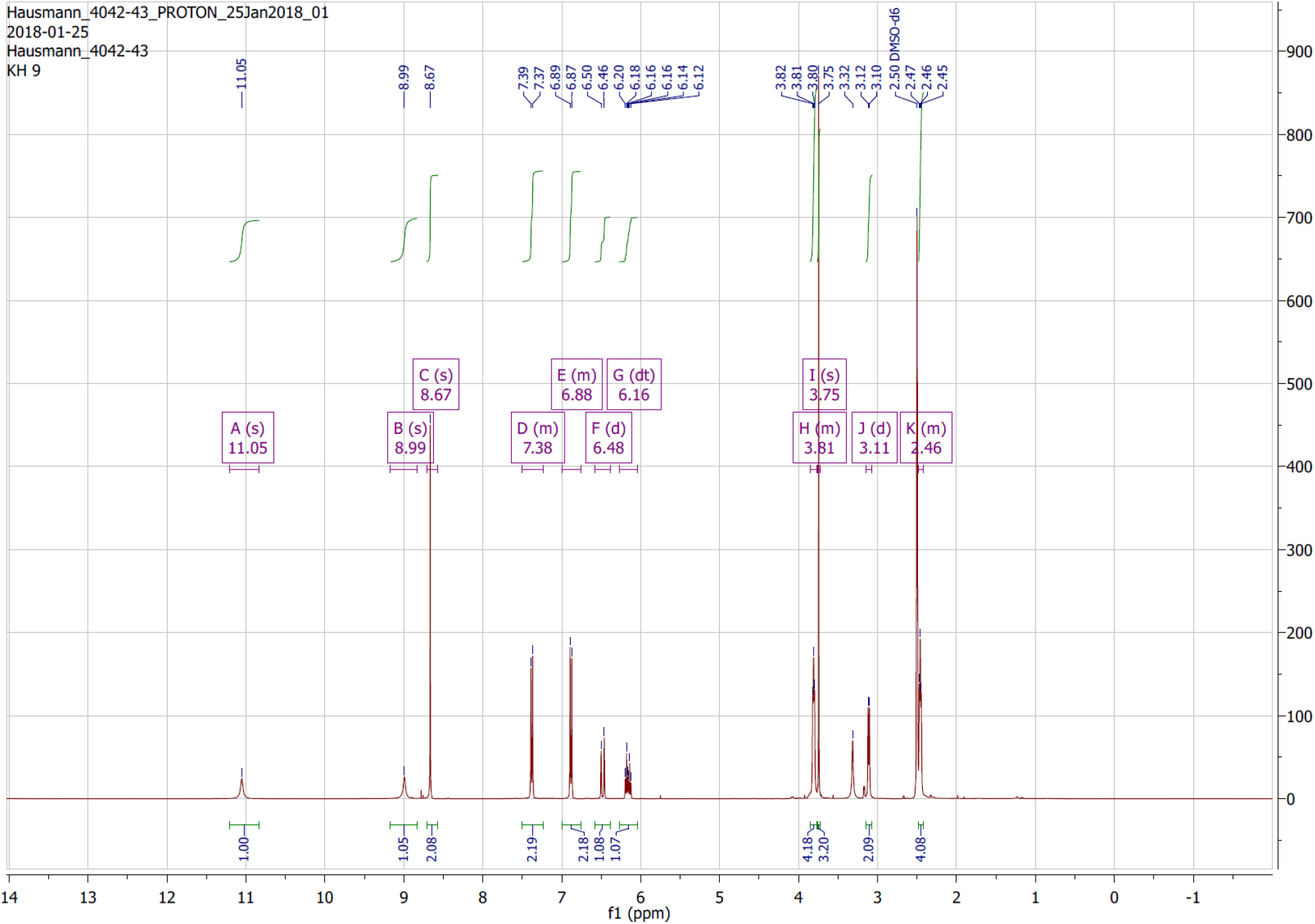
^1^H NMR spectra KH9.

**Figure S 2.**
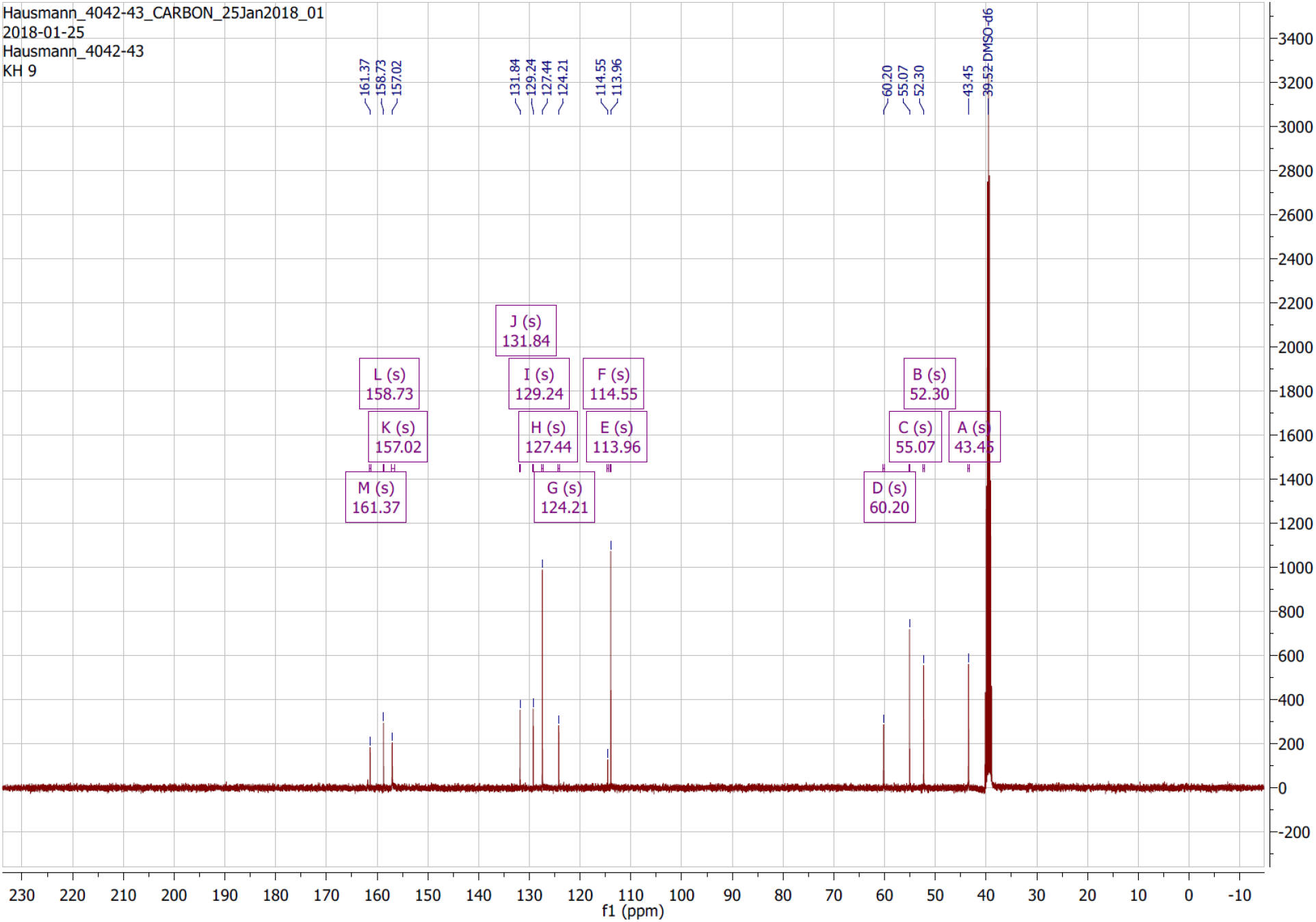
^13^C NMR spectra KH9.

**Figure S 3.**
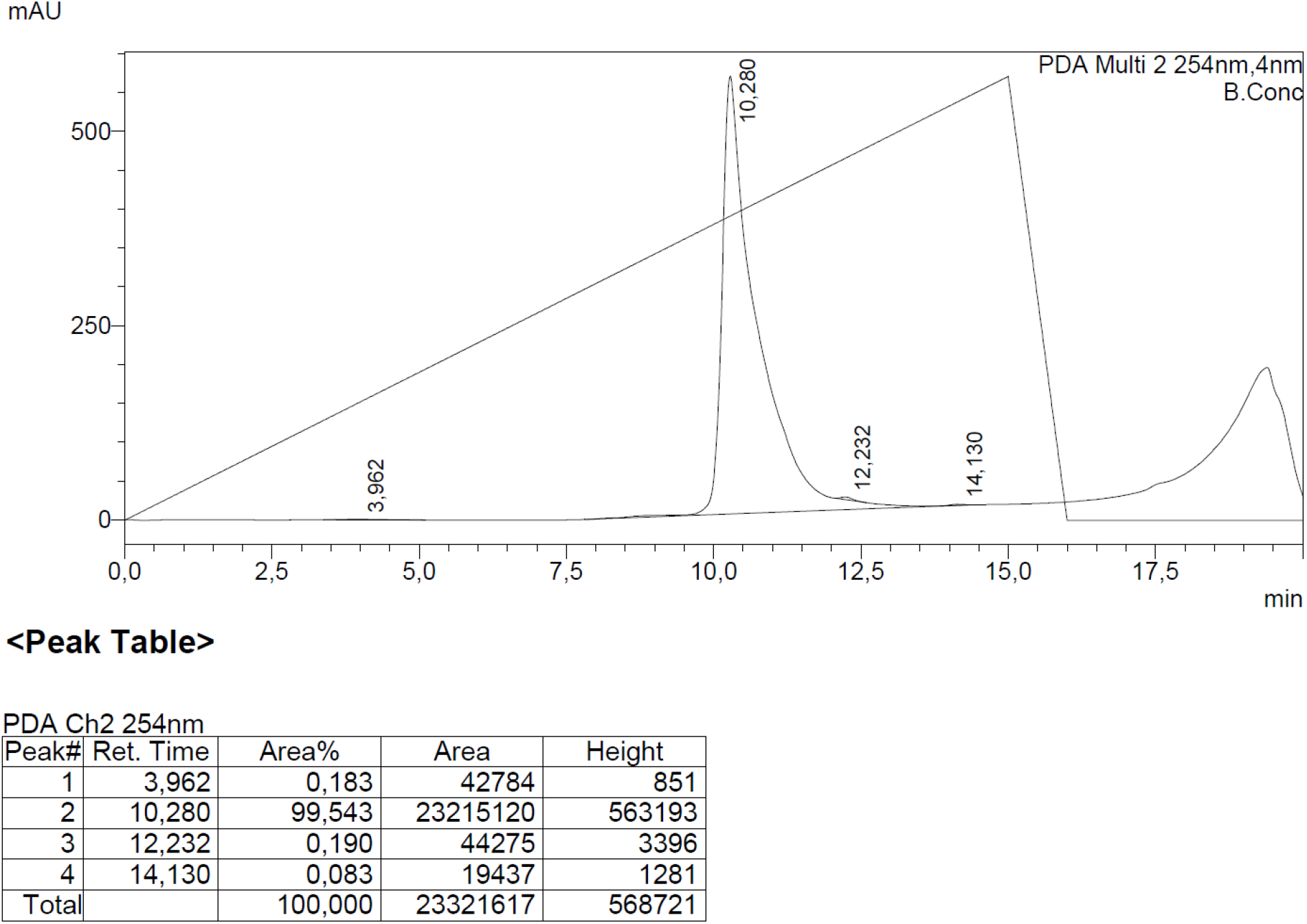
HPLC KH9

**Figure S 4.**
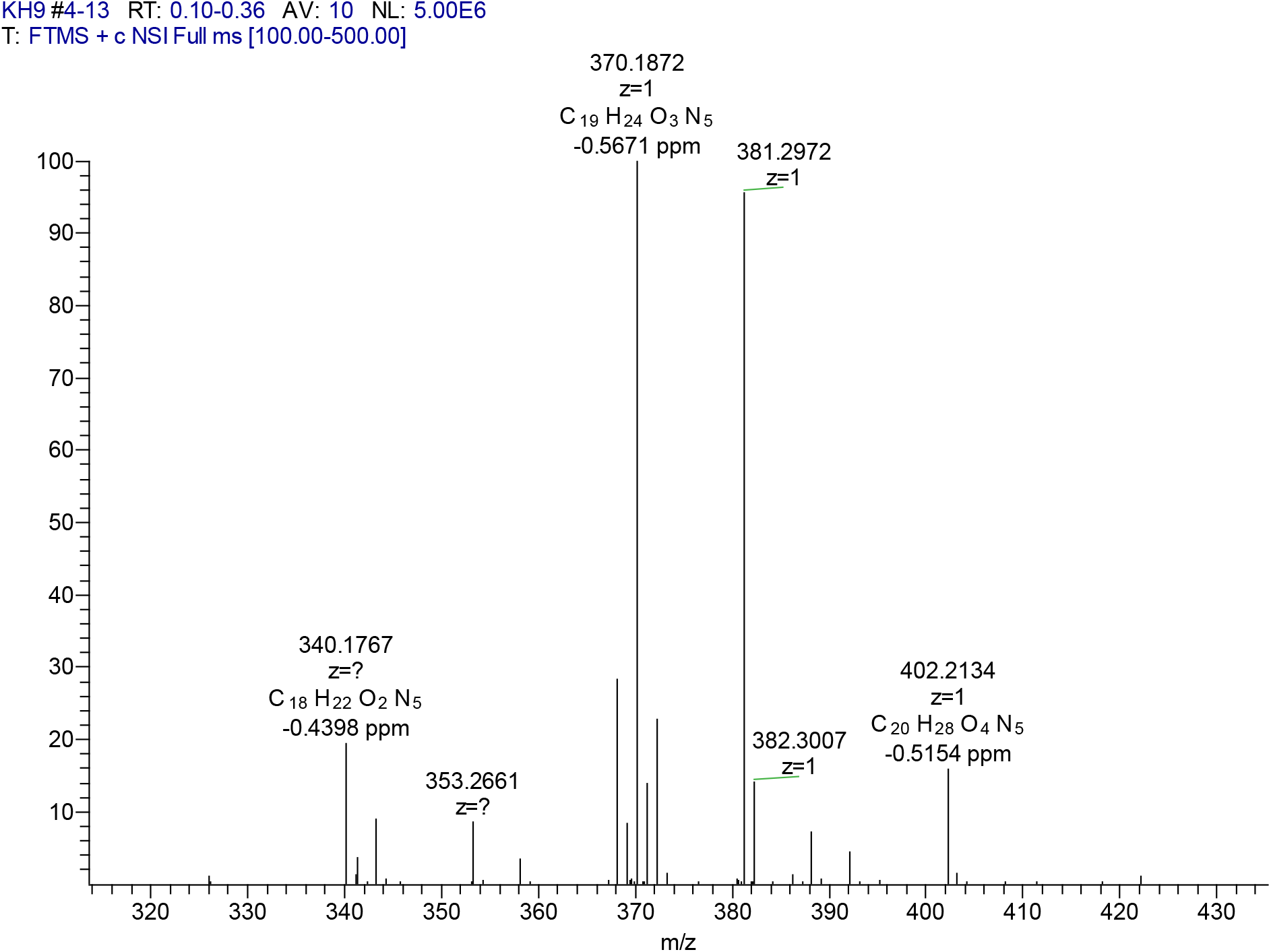
HRMS KH9.

### KH16 (7b)

**Figure S 5.**
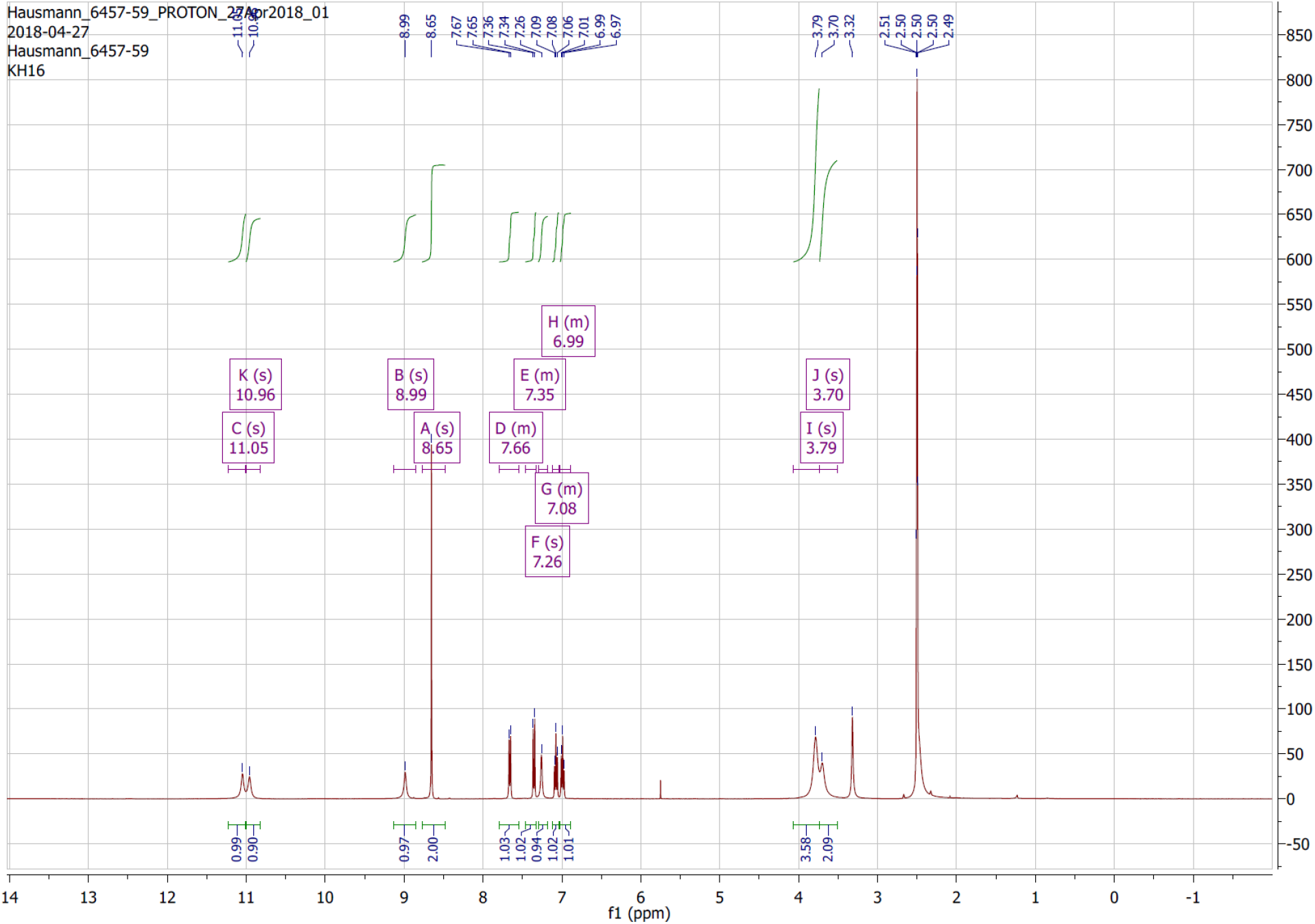
^1^H NMR spectra KH16.

**Figure S 6.**
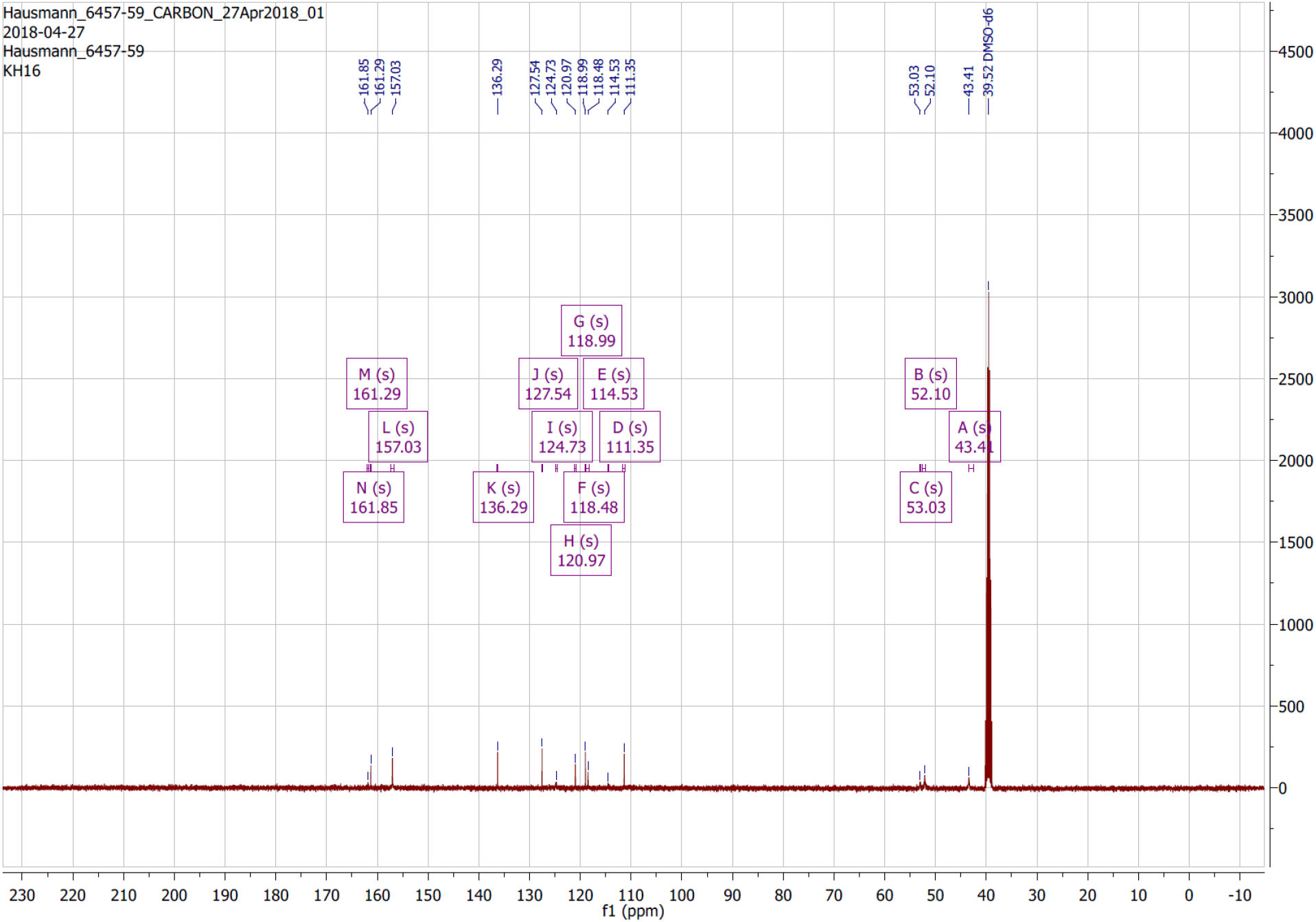
^13^C NMR spectra KH16.

**Figure S 7.**
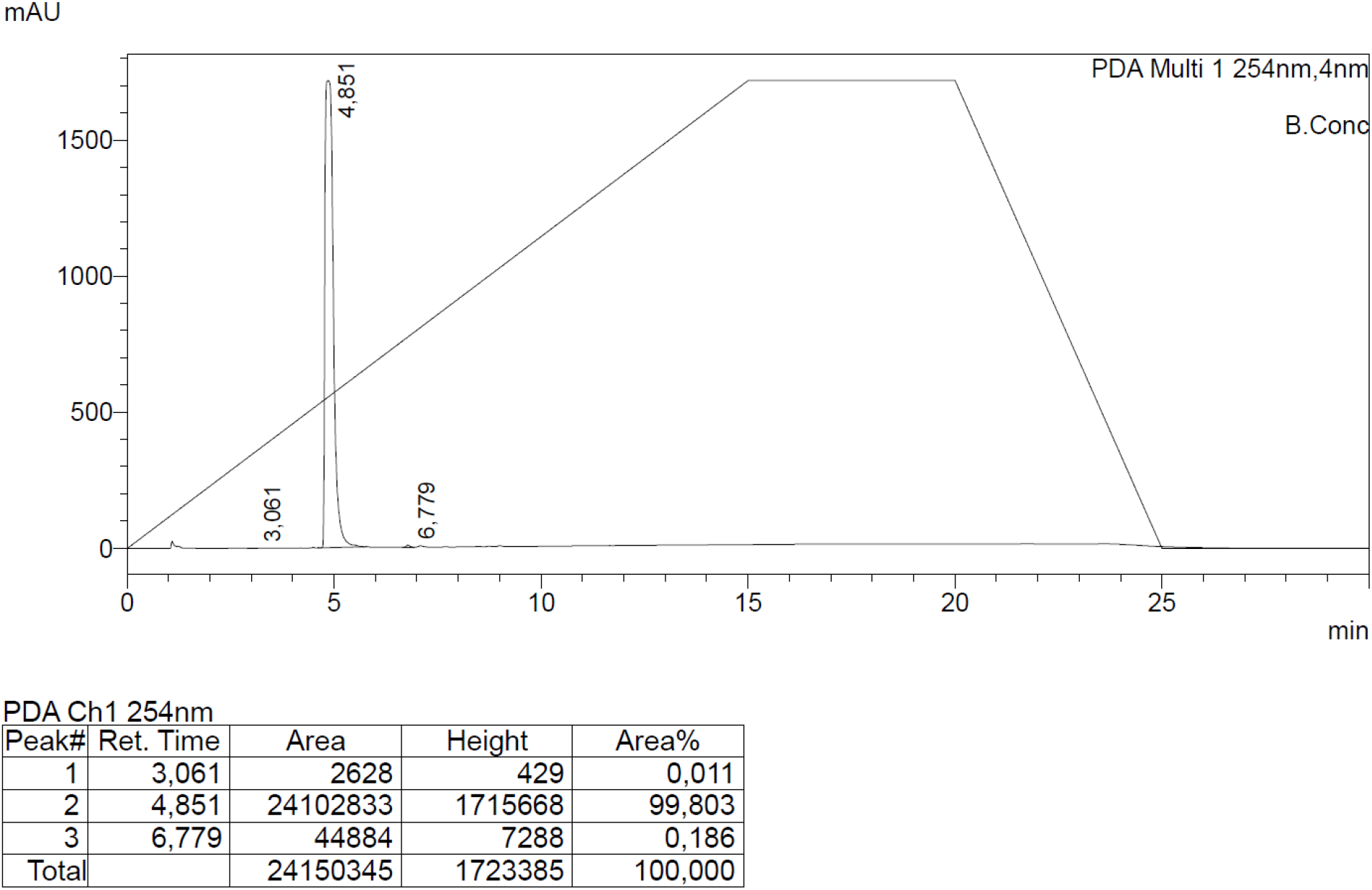
HPLC KH16.

**Figure S 8.**
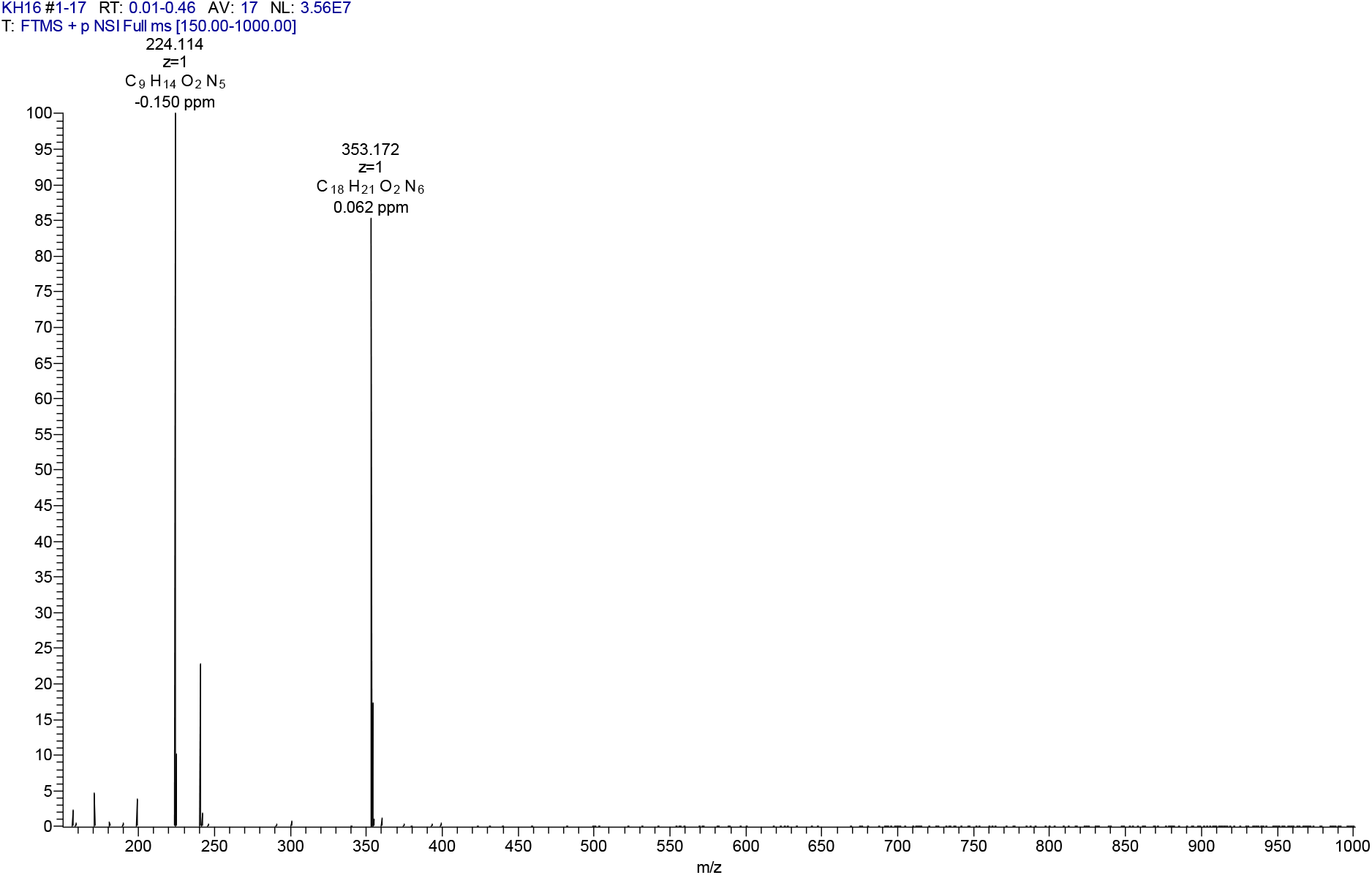
HRMS KH16.

### KH29 (7c)

**Figure S 9.**
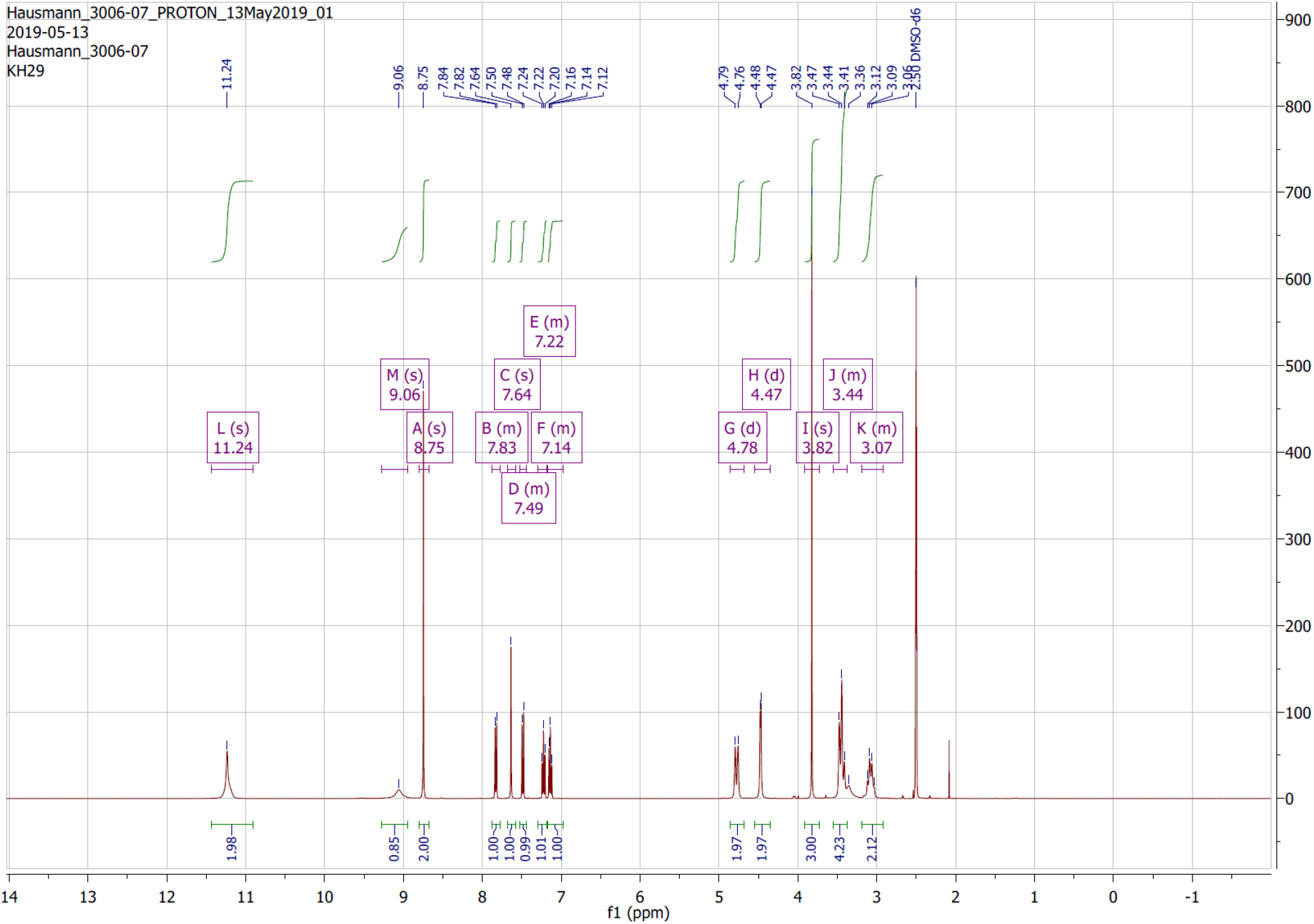
^1^H NMR spectra KH29.

**Figure S 10.**
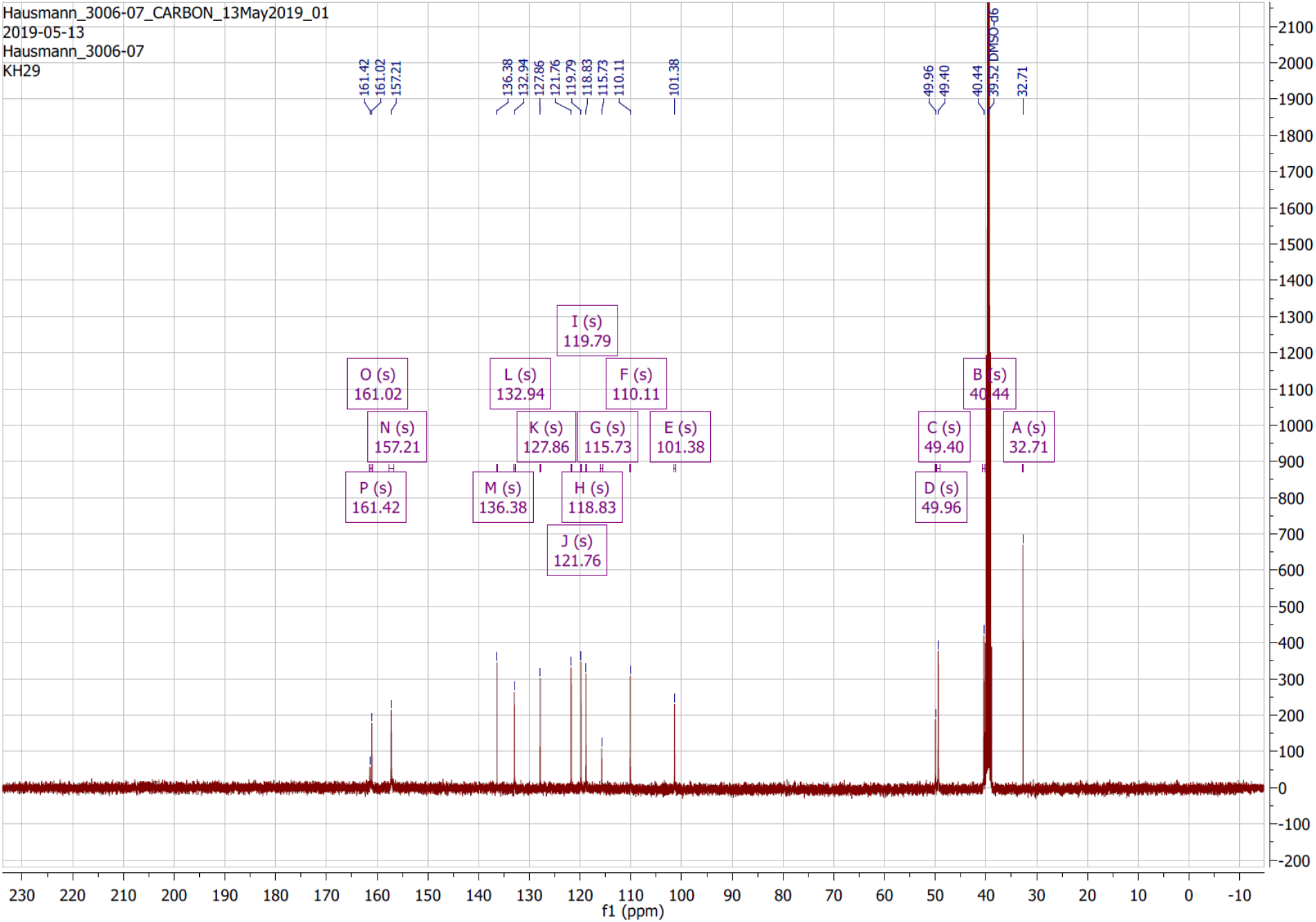
^13^C NMR spectra KH29.

**Figure S 11.**
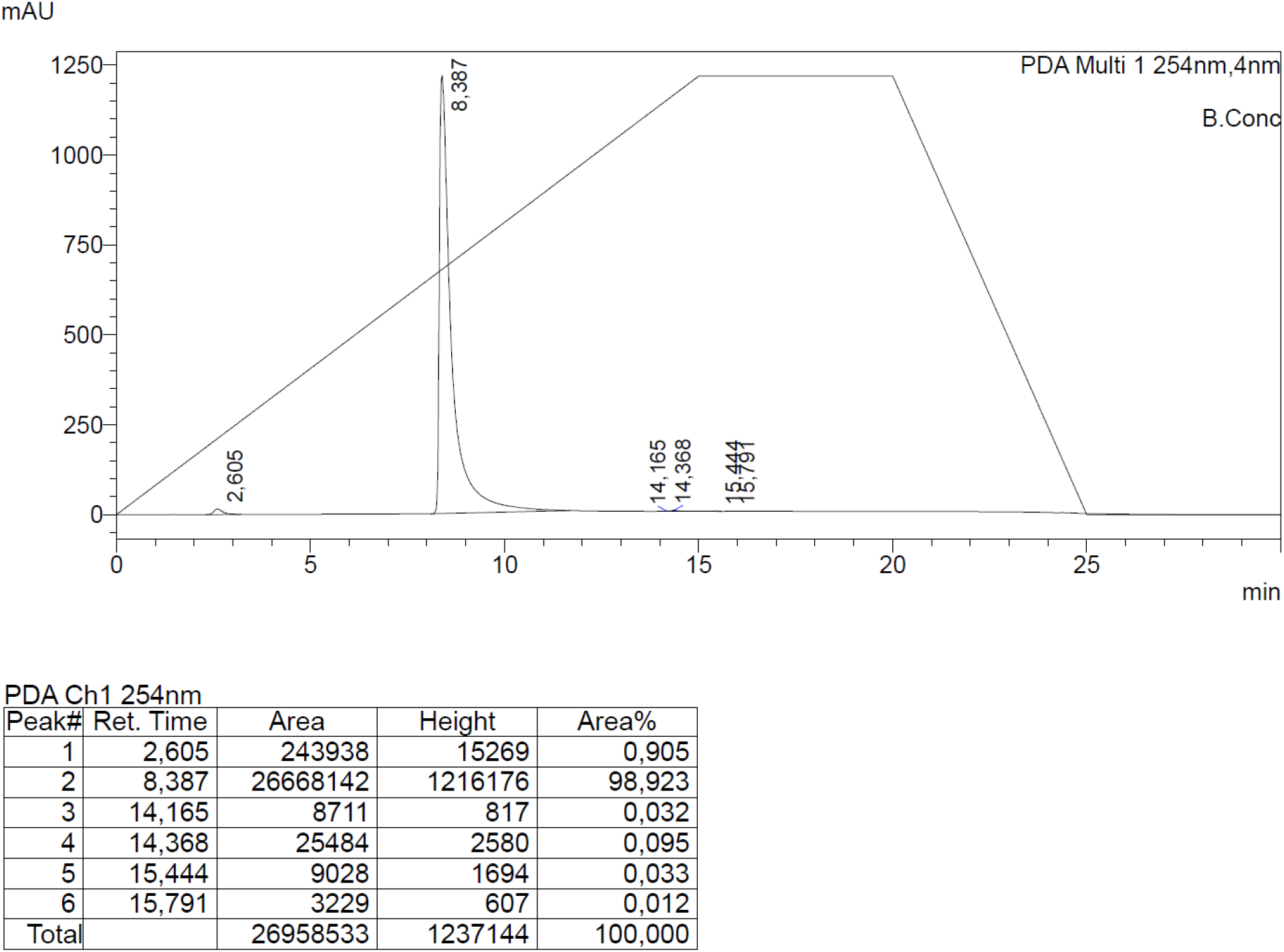
HPLC KH29

**Figure S 12.**
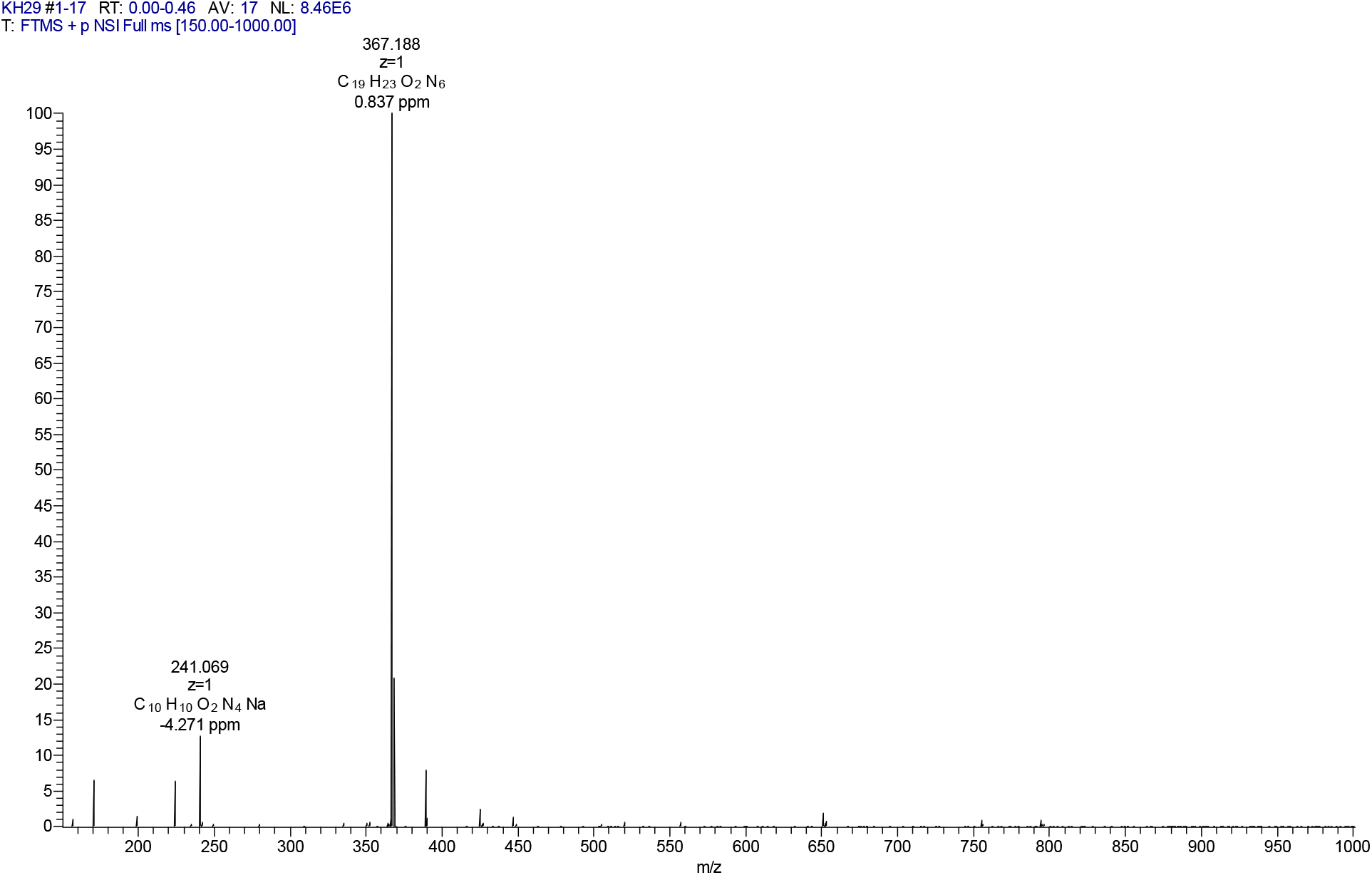
HRMS KH29

### In vitro HDAC inhibition – IC_50_ plots

**Fig S13.**
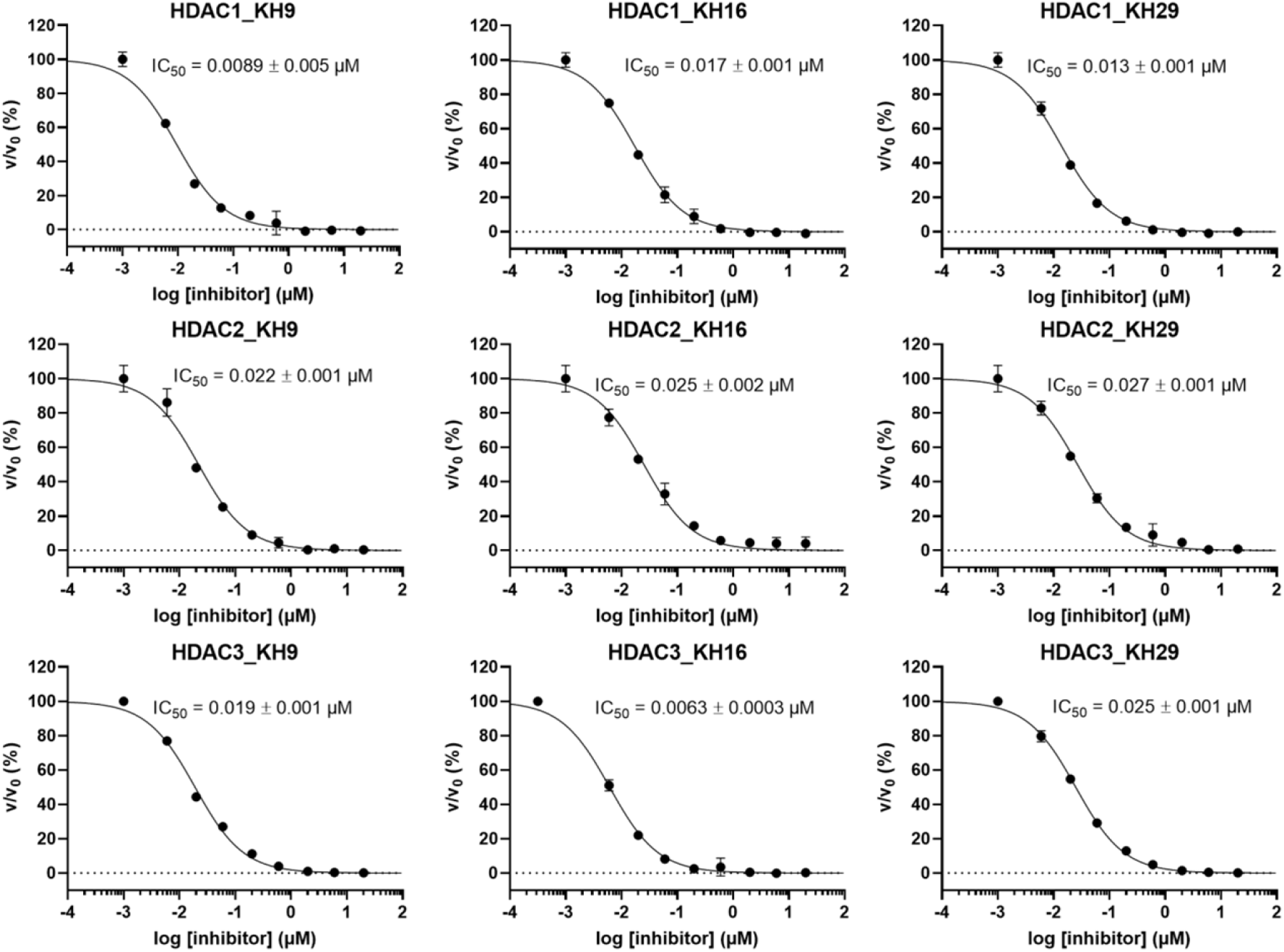
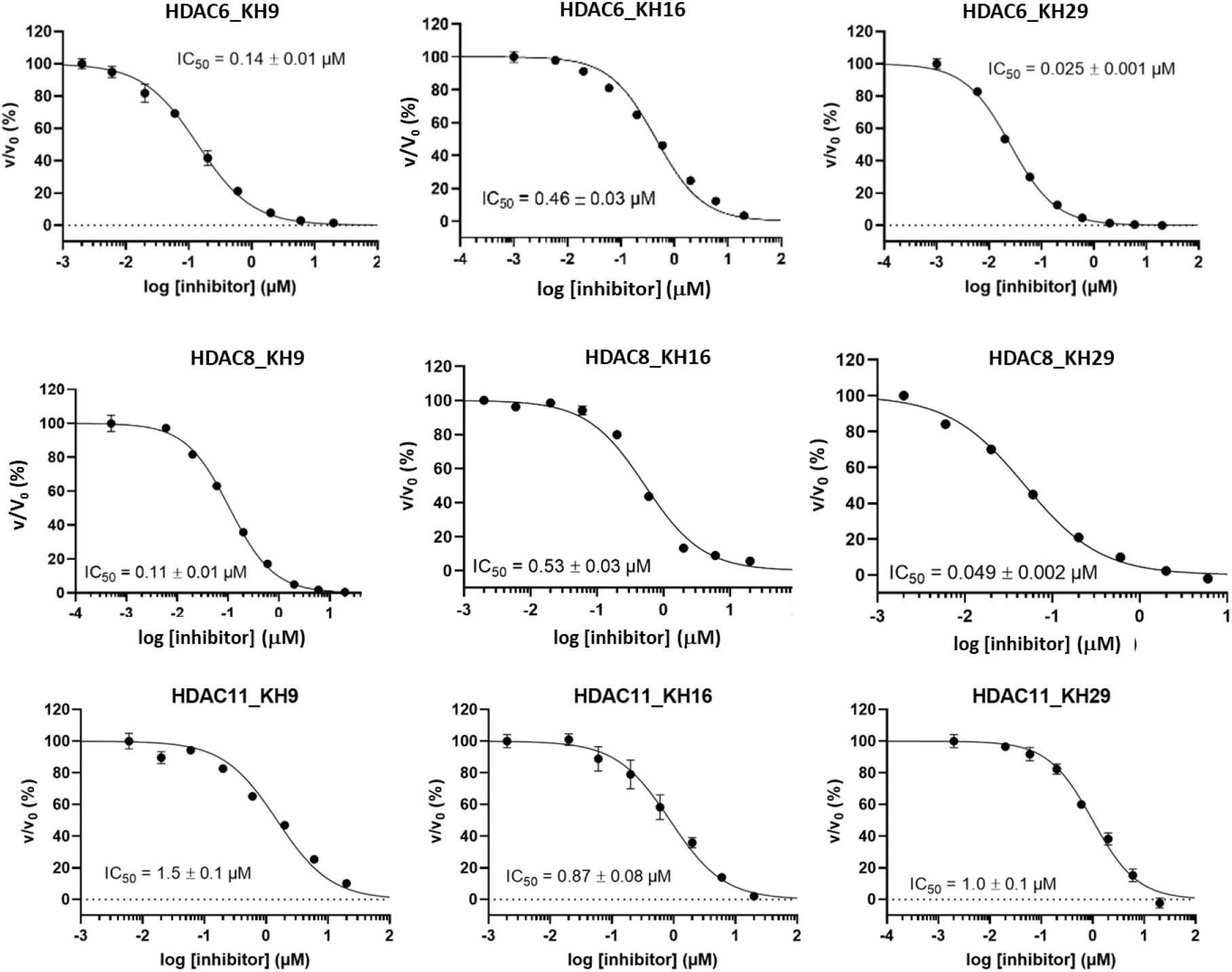
IC_50_ curves of KH9, KH16, KH29 for tested HDACs

